# Lymphatics unload VEGFR3 in response to the alteration of VEGFR2-mediated transcriptional programs

**DOI:** 10.1101/2025.10.20.683599

**Authors:** Taotao Li, Xudong Cao, Fei Zhou, Xiujuan Li, Wenjuan Ma, Xiuyan Zhang, Haoyu Zhong, Beibei Xu, Man Chu, Xiwen Jia, Kai Ding, Xin Shen, Yahui Liu, Yun Zhao, Zhen Zhang, Junhao Hu, Young-Kwon Hong, Lena Claesson-Welsh, Yulong He

## Abstract

Vascular endothelial growth factor receptor 2 (VEGFR2) is a key target for regulating the endothelial cell lineage and angiogenesis. It is also expressed by lymphatic endothelial cells (LEC) while its participation in the process of lymphangiogenesis remains inadequately characterized. We show in this study that VEGFR2 is highly expressed in dermal initial lymphatic vessels and valves. The induced deletion of pan-endothelial *Vegfr2* at the neonatal stage produced a potent suppression of dermal lymphatic growth, including the thinner lymphatic diameter, the decrease of LEC number and lymphatic valves. Mechanistically, the VEGFR2 insufficiency led to a dramatic decrease of lymphatic VEGFR3, a key regulator mediating signals for lymphatic growth and remodeling. The RNA sequencing analysis revealed that GO terms enriched for the downregulated genes included biological processes related to the EC development while pathways related to the hematopoiesis and immune responses were upregulated in the skin of *Vegfr2* mutants compared with the littermate controls. This was further confirmed by the RNA-seq analysis of dermal tissues 48 hours after the endothelial *Vegfr2* deletion. Consistently, an inhibitory effect on the dermal lymphatic growth was also observed by targeting *Vegfr2* in PROX1^+^ cells, manifesting a similarly altered transcriptome signature. The alteration of lymphatic gene expression was further validated by the siRNA mediated knockdown of *Vegfr2* in primary lymphatic endothelial cells, showing a transcriptional trend of LEC to hematopoietic transition. Findings from this study imply that VEGFR2 is required for the maintenance of endothelial identity and its insufficiency may trigger the alteration of lymphatic transcriptional programs to diminish the VEGFR3-mediated lymphangiogenesis.

## Introduction

The endothelial cell (EC) differentiation from the mesoderm is crucial for the establishment of blood vascular and lymphatic systems during the embryogenesis in mammals. The molecular programs governing the EC lineage commitment have been extensively investigated, leading to the identification of a number of key factors and pathways in the process. Vascular endothelial growth factor receptor 2 (VEGFR2) mediates a principal pathway for the vascular development. It has been shown that VEGFR2 is a direct target of MECOM (MDS1 And EVI1 Complex Locus), a leading factor in the regulation of the transcriptional program for the EC differentiation [1]. PROX1 (a homeobox prospero-like protein) is required for the determination and maintenance of lymphatic endothelial cell (LEC) identity and regulates the expression of VEGFR3 [2–6]. Lymphatic formation and remodeling are governed by a plethora of organotypic lymph/angiogenic pathways [7–11]. Among them, vascular endothelial growth factor family members (VEGFs) and their receptors are the most prominent regulators [12–17]. VEGFC/VEGFR3-mediated pathways are crucial for the formation of lymphatic networks and insufficient VEGFR3 signals lead to lymphatic dysplasia and lymphedema [16–20]. Lymphatic endothelial cells also express VEGFR2 [21, 22], a key target of the transcription factor MECOM for the determination of endothelial cell lineage [1]. The fully processed active forms of VEGFC and VEGFD can activate VEGFR2 or exert biological functions via the VEGFR2/VEGFR3 heterodimers [22–24]. The soluble form of VEGFR2 was shown to be an endogenous inhibitor of lymphangiogenesis in the cornea [25]. In zebrafish, VEGFR2 was reported to regulate lymphatic development in an organotypic manner [26]. Deletion of *Vegfr2* using the *Lyve1-Cre* transgenic mice was shown to decrease the dermal lymphatic vessel density, but with the normal lymphatic diameter and the draining function [27]. On the other hand, the recombinant adenovirus-mediated overexpression of VEGFA or VEGFE (a VEGFR2-specific factor) induced abnormal lymphangiogenesis [28, 29]. It is worth noting that over-activation of VEGFR2 by excess VEGFA could induce blood vascular leakage and in turn affect the integrity of lymphatic system. Suppression of the VEGFA/VEGFR2-induced vascular leakage and interstitial edema may be responsible for the inhibition of lymphangiogenesis and lymphatic dissemination as shown in the *Vegfr2^Y949F/Y949F^* mice [30].

Although VEGFR2 has been shown to participate in the regulation of lymphatic vessel growth, the underlying mechanism is largely unknown. To further investigate its function in LECs, we employed genetic mouse models targeting *Vegfr2* in either pan-endothelial cells (*Vegfr2^iKO;CDH5+^*) or in PROX1^+^ endothelial cells (*Vegfr2^iKO;PROX1+^*) [31, 32]. We show in this study that the induced deletion of pan-endothelial or lymphatic *Vegfr2* suppressed lymphatic growth. We demonstrate that VEGFR3 was dramatically reduced in dermal lymphatic vessels upon the VEGFR2 insufficiency. The alteration of VEGFR3 was further validated by the siRNA mediated knockdown of *Vegfr2* in primary lymphatic endothelial cells. Gene ontology analysis revealed that pathways related to hematopoiesis and immune responses (involving INFα/γ and NFκB) were enriched in the upregulated dermal genes after the induced deletion of endothelial or lymphatic *Vegfr2*, or by the siRNA-mediated knockdown of *Vegfr2* in the primary lymphatic endothelial cells (LECs). It has been shown recently that lymphatic endothelial cells have hemogenic capacity which is repressed by the lymphatic fate determinant PROX1 [33]. Findings from this study imply that VEGFR2 is required for the maintenance of postnatal endothelial cell identity and its insufficiency may lead to a transcriptional trend of LEC to hematopoietic transition to diminish the VEGFR3-mediated lymphangiogenesis.

## Materials and Methods

### Mouse models

All animal experiments were approved by the Animal Care Committee of Soochow and Nanjing University Animal Center (MARC-AP#YH2 /SUDA20250507A03). All the mice used in this study were housed in a SPF (specific pathogen free) animal facility with a 12/12 hours dark / light cycle, and were free to food and water access. Normal mouse diet (Suzhou Shuangshi Experimental Animal Feed Technology Co., Ltd.) and cage bedding (Suzhou Baitai Laboratory Equipment Co., Ltd.) were used. To generate mice with cell-specific *Vegfr2* gene deletion, *Vegfr2^Flox/Flox^* mice [34] were crossed with transgenic mice expressing Cre recombinase in pan-endothelial cells (*Cdh5-CreERT2*) [31] or lymphatic endothelial cells (*Prox1-CreERT2*) [32].

The floxed mice used in this study were maintained on the C57BL/6J background with at least five backcrosses. *Vegfr2^Flox/Flox^;Rosa26^RFP/RFP^* mice was generated as previously reported and expressed the red-fluorescence protein tdTomato after Cre-mediated recombination [35]. *Rosa26-mTmG* mice were purchased from Model Animal Research Center of Nanjing University. The *Cdh5-CreERT2;Rosa26-mTmG* and *Prox1-CreERT2;Rosa26-mTmG* lines were generated by two rounds of crossing. *Prox1-EGFP* mice were generated as previously reported [36]. In all the phenotype analysis, littermates were used as control. Mice were sacrificed by asphyxiation with rising concentration of carbon dioxide gas, followed by cervical dislocation and tissue collection.

### Induction of gene deletion

Induction of gene deletion was performed by the systemic administration of tamoxifen (Sigma-Aldrich, T5648). Briefly, new-born pups, including the *Vegfr2^Flox/Flox^*;*Rosa26^RFP/RFP^*;*Cdh5-CreERT2* (named *Vegfr2^iKO;CDH5+^*), *Vegfr2^Flox/Flox^ or Vegfr2^Flox/-^;Rosa26^RFP/RFP^*;*Prox1-CreERT2* (named *Vegfr2^iKO;PROX1+^*) and littermate control mice (*Vegfr2^Flox/Flox^;Rosa26^RFP/RFP^*or *Vegfr2^Flox/+^*;*Rosa26^RFP/RFP^*) were treated with tamoxifen (60 μg per day) by four daily intragastric injections (postnatal days 1-4, P1-4) or treated with tamoxifen (200 μg per day) by three or five daily intraperitoneal injections (P10-12/14,). For the *Prox1-CreERT2;Rosa26-mTmG* mice, tamoxifen treatment was performed at P1-4 by intragastric injections (60 μg per day). Mice were analyzed at different time points as detailed in each experiment.

### RNA sequencing analysis

Ear skin (P21) or abdominal skin (P3) tissues were harvested from control and *Vegfr2* mutant mice with the induced gene deletion in endothelial cells as described above. Tissues were kept frozen in liquid nitrogen, and total RNA was extracted from the tissues using Trizol (Invitrogen) according to the manufacturer’s instruction. mRNA was purified for the preparation of RNA-seq libraries as previously described [37]. Sequencing was performed on DNBSEQ platform with PE150 (read length, BGI-Shenzhen, China). The sequencing data were processed according to the predefined analysis pipeline. The differential expression analysis was conducted using the R package DESeq2 (v 1.42.1). Functional enrichment analysis was performed using clusterProfiler (v4.10.1). The hallmark gene sets used in the GSEA (Gene Set Enrichment Analysis) are mainly defined according to the transcriptome atlas of murine endothelial cells, the molecular signatures database (mh.all.v2024.1.Mm.symbols) and the related GO terms [37–41]. The raw and processed data of RNA-seq analysis will be deposited in GEO.

### Immunostaining

For the whole-mount immunostaining, tissues were harvested and processed as previously described [42]. Briefly, the tissues were fixed in 4 % paraformaldehyde (PFA, Sigma-Aldrich, 158127), blocked with 3 % (w/v) milk (Valio, skimmed milk powder instant) in PBS-TX (0.3 % Triton X-100, VETEC, V900502), and incubated with primary antibodies and the corresponding secondary antibodies overnight at 4 °C. For the immunofluorescence staining of the cultured lymphatic endothelial cells, cells were grown on coverslips placed at the bottom of a culture dish and coated with 0.2% gelatin (Sigma-Aldrich, G2500). Cells were fixed in 4% PFA for 15 min, washed with PBS, permeabilized with 0.1% PBS-TX for 20 min and blocked with 5% fetal calf serum (FCS, PromoCell, C-37360) in PBS for 60 min at room temperature (RT). Cells were then incubated with primary antibodies overnight at 4°C, washed with PBS followed by the incubation with secondary fluorescently conjugated antibodies for 2 hours at RT. The primary antibodies used were anti-PECAM1 (BD Pharmingen, 553370, 1:500), anti-LYVE1 (Abcam, ab14917, 1:1000), anti-PROX1 (R&D, AF2727, 1:500), anti-PROX1 (ReliaTech, 102-PA32S, 1:200), anti-VEGFR2 (R&D, AF644, 1:300), anti-VEGFR3 (R&D, AF743, 1:200), anti-CD144 (BD Pharmingen, 550548, 1:300), anti-ITGA9 (R&D, AF3827, 1:200), anti-CD45 (BD Pharmingen, 550539, 1:500) and anti-DLL4 (R&D, AF1389, 1:300). The secondary antibodies used were Alexa488 Fluor-conjugated (Invitrogen, A21206 and A21208, 1:500), Alexa568 Fluor-conjugated (Invitrogen, A78946), Cy3-conjugated (Jackson ImmunoResearch Labs, 705-165-147, 1:500) and Cy5-conjugated (Jackson ImmunoResearch Labs, 712-175-153, 711-175-152 and 705-175-147, 1:500). Samples were mounted with 50 % glycerol (Sigma-Aldrich, G5516) containing DAPI (Beyotime, C1002) and analyzed with the Olympus FluoView 3000 confocal microscope.

### Isolation of lymphatic endothelial cells

Isolation of primary lymphatic endothelial cells was performed following the protocol by Kazenwadel et al. [43]. Briefly, the mouse line carrying the CAG-bgeo-tsA58T Ag transgene was crossed with *Tek-Cre* mice to turn on the expression of tsA58T Ag in endothelial cells [44, 45]. The transgenic mice (6-10 week-old) were lethally anesthetized using 0.8% sodium pentobarbital by intraperitoneal injection and perfused with PBS followed by the digestion mix in the DMEM /20% FCS supplemented with collagenase type II (2.5 mg/ml; Sigma C6885) and type Ia (2.5 mg/ml; Sigma C2674). Lung tissues were collected, minced and digested in the digestion mix at 37 °C for 30 min. The digested cell mix was passed through 40 μm cell strainer (BD Falcon, 352340) and then rinsed with ice-cold wash buffer. The resuspended cells were separated using the miniMACS Separation System (Miltenyi Biotech, 130-042-201), together with the goat anti-rabbit IgG microbeads (Miltenyi Biotech, 130-048-602) or goat anti-rat IgG microbeads (Miltenyi Biotech, 130-048-501). Hematopoietic cells were depleted from the cell mixture using miniMACS and rat anti-mouse F4/80 (SeroTech, MCAP497) and CD45 antibodies (BD Pharmingen, 550539). The remaining cells were selected for endothelial cells (ECs) by miniMACS and anti-mouse PECAM1 (BD Pharmingen, 553370) and then for lymphatic endothelial cells by anti-mouse LYVE1 (Abcam, 14917). The isolated primary lymphatic endothelial cells were cultured at 33 °C in the EC medium (ScienCell, 1001).

### siRNA transfection

To explore the functional mechanism of VEGFR2 in lymphatic endothelial cells, murine primary lymphatic endothelial cells (LEC) were isolated as described above and cultured to approximately 70-80% confluence. LECs were transfected with siRNA duplex oligoribonucleotides targeting mouse *Vegfr2* using CALNP™ RNAi in vitro transfection reagent (DN001) according to the manufacturer’s instruction. The knockdown efficiency was examined by the western blotting analysis 48 hours after the transfection. RNA from the transfected and control cells was also extracted for the RNA-seq analysis. Two siRNAs for mouse *Vegfr2* were used (siRNA-#1: sense, 5′-CCCGUAUGCUUGUAAAGAAdTdT-3′; antisense, 5′-UUCUUUACAAGCAUACGGGdCdT-3′; siRNA-#2: sense, 5′-CCGAAUCCCUGUGAAGUAUdTdT-3′; antisense, 5′-AUACUUCACAGGGAUUCGGdAdC-3′). The control scrambled siRNA (sense, 5′-UUCUCCGAACGUGUCACGUdTdT-3′; antisense, 5′-ACGUGACACGUUCGGAGAAdTdT-3′) was used as a negative control.

### Western blotting

Mouse skin tissues were harvested, snap-frozen in liquid nitrogen and stored at −80 °C freezer. Tissues or isolated cells were lysed in NP-40 lysis buffer (Beyotime, P0013F) supplemented with protease inhibitor cocktail (complete Mini, Roche 04693124001), phosphatase inhibitor cocktail (PhosSTOP, Roche, 04906837001) and 1 mmol/L PMSF (Bio Basic, PB0425). Protein concentration was determined using the BCA protein assay kit (PIERCE, 23227), and equal amounts of protein were used for analysis. Briefly, after the protein transfer from gels to PVDF membranes (Millipore, IPVH00010) and the antibody incubations, images were acquired by the chemiluminescent detection method (PerkinElmer, NEL105001EA) using X-ray film (XBT, Carestream, 6535876) or Touch Imager (eBlot). For images acquired using X-ray film, the protein markers were manually marked on the films overlapped with the PVDF membranes containing the prestained protein ladder (ThermoFisher Scientifc, 26619). The blots were washed and re-probed with antibodies for beta-Actin as loading controls. The antibodies used in this study include anti-VEGFR3 (R&D, AF743, 1:1000), anti-PROX1 (R&D, AF2727, 1:1000), anti-VEGFR2 (R&D, AF644, 1:1000), anti-beta-Actin (Santa Cruz, sc-47778, 1:20000) and Bovine-anti-Goat IgG, HRP Conjugated (Jackson ImmunoResearch Labs, 805-035-180, 1:5000).

### Flow cytometry and peripheral blood analysis

Cells from mouse bone marrow were harvested in 2% (vol/vol) fetal bovine serum-supplemented Hank’s Balanced Salt Solution (collectively called HF). The cells were blocked with 2% HF containing CD16/32 (Becton Dickinson, 223142), and stained with antibodies including anti-CD4 (Biolegend, 100510, 1:100), anti-CD8 (Biolegend, 100712, 1:100), anti-B220 (Biolegend, 103205, 1:100), anti-CD11b (BD Pharmingen, 557657, 1:100), anti-Gr1 (Biolegend, 108406, 1:100) and anti-TER-119 (eBioscience, 47-5921-82, 1:100) for flow cytometry (Gallios). For the analysis of peripheral blood cells, mice were anaesthetized using 0.8% sodium pentobarbital by intraperitoneal injection and blood samples were collected from the retro-orbital plexus in 1.5 ml microcentrifuge tubes coated with EDTA-K2 (SKJYLEAN, SK-3542). Blood cell counts were obtained using an automated blood analyzer (URIT-5160Vet), including red blood cells, white blood cells, neutrophils, lymphocytes, monocytes, eosinophils, basophils and platelets.

### Statistical analysis

For the 2-group comparison, the unpaired t test was performed with Welch correction if data passed the D’Agostino-Pearson normality test, or the unpaired nonparametric Mann-Whitney U test was applied using GraphPad Prism v8.0. Data are expressed as mean ± SD. All statistical tests were 2-sided.

## Results

### Thinner lymphatic diameter after the endothelial deletion of *Vegfr2*

We found that VEGFR2 was differentially expressed in lymphatic vessels among tissues. It was highly expressed in the initial lymphatic vessels of dermal tissues, but relatively low in lymphatics of several other tissues examined including the intestinal villi and trachea (Supplemental Fig. 1). To investigate the role of VEGFR2 in lymphatic development, we employed the inducible pan-endothelial Cre deleter (*Cdh5-CreERT2*) for the generation of endothelial *Vegfr2* deleted mice (*Vegfr2^iKO;CDH5+^*). The gene deletion was induced by the tamoxifen treatment via the intragastric injection at postnatal days 1 to 4 (P1-4), and the *Vegfr2^iKO;CDH5+^*and littermate control mice were analyzed at P7 (Supplemental Fig. 2A). The expression of reporter RFP indicated the Cre-mediated recombination in endothelial cells (ECs). The efficiency of *Vegfr2* deletion was also confirmed by the immunostaining of lymphatics for VEGFR2 together with LYVE1 (Supplemental Fig. 2C). The induced deletion of endothelial *Vegfr2* resulted in a dramatic suppression of angiogenesis as observed in the retina, skin, intestinal villi and trachea (Supplemental Fig. 2B-D, F and H). In contrast, the lymphatic phenotypes differed among tissues in the *Vegfr2^iKO;CDH5+^* mice, with a significant suppression of lymphangiogenesis in the skin but not in the intestinal villi and trachea (Supplemental Fig. 2C and E-H).

To minimize the effects on the blood vasculature as a consequence of the endothelial *Vegfr2* deletion, we performed the tamoxifen treatment from postnatal day 10-12 (P10-12, *Vegfr2^iKO;CDH5+^*) and mice were analyzed at P21 (Fig. 1A). Immunostaining analysis of the ear skin showed that the blood vascular network appeared normal in *Vegfr2* mutant mice. The slight decrease of PECAM1^+^ vascular density in *Vegfr2^iKO;CDH5+^* mice could partly result from the decrease of lymphatic vessels (Fig. 1B-D). In line with the results described above, we found that the loss of endothelial VEGFR2 resulted in a dramatic decrease in the lymphatic diameter and LYVE1^+^ vessel area (Fig. 1B-C and E-F). Consistently, there was a significant decrease in the number of lymphatic endothelial cells in the initial as well as the collecting lymphatic vessels (Fig. 1G-I). Furthermore, the endothelial VEGFR2 insufficiency also led to a significant decrease in the lymphatic valves of pre-collectors (Fig. 1G and J), suggesting the aberrant lymphatic remodeling and maturation.

**Fig. 1.**
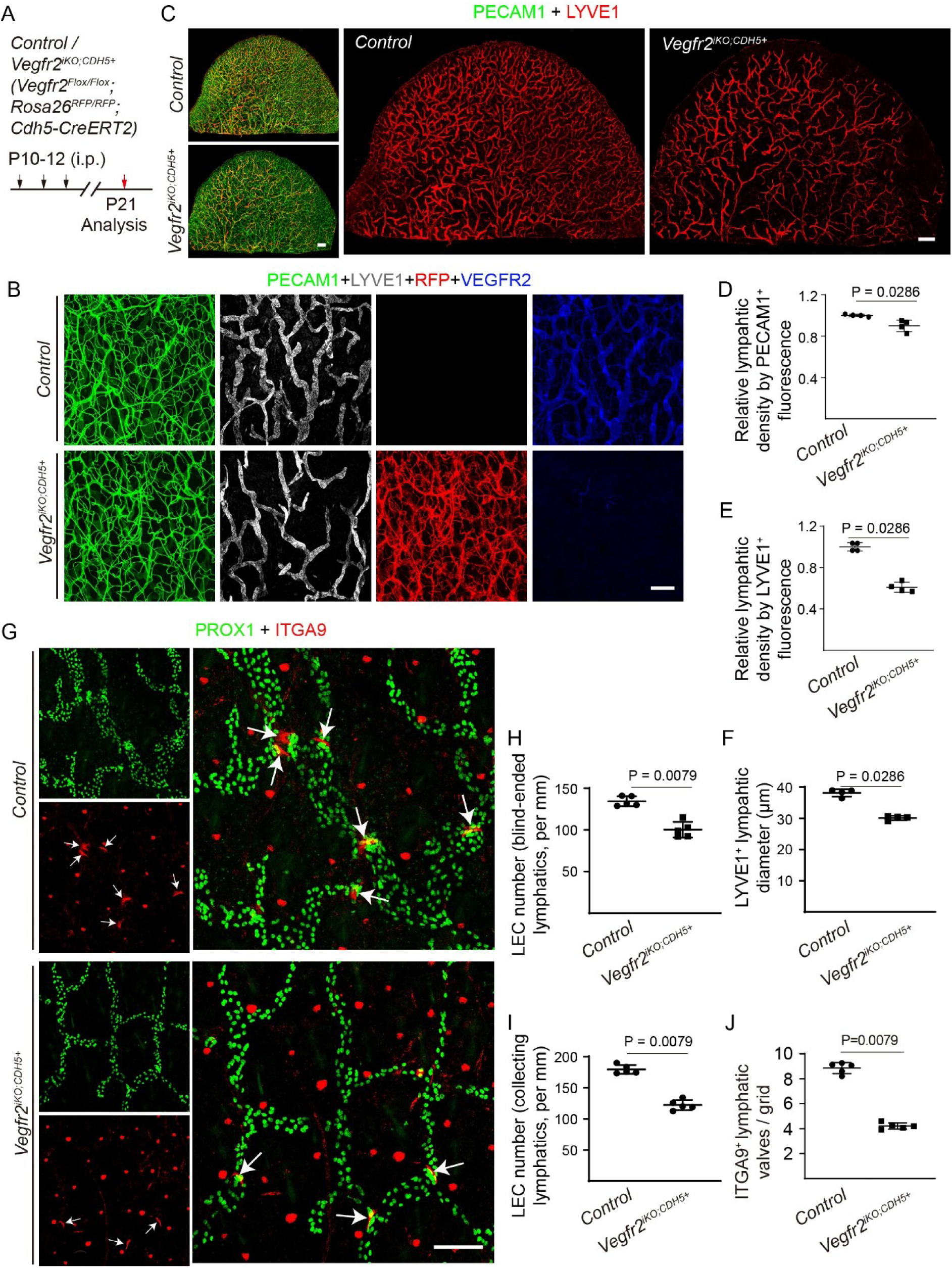
A phenotype of decreased lymphatic diameter and LEC numbers by targeting *Vegfr2* in CDH5^+^ ECs. **A.** Tamoxifen intraperitoneal administration (i.p.) and mice analysis scheme. **B.** Analysis of VEGFR2 deletion by the whole-mount immunostaining of blood and lymphatic vessels in the ventral side of ear skin from the *Vegfr2^iKO;CDH5+^* and control mice (P21) for PECAM1 (green), LYVE1 (white) and VEGFR2 (blue). The reporter RFP signals indicate the Cre (*Cdh5-CreERT2*)-mediated recombination in blood and lymphatic vessels of the *Vegfr2^iKO;CDH5+^*mice. **C.** Analysis of ear skins from the *Vegfr2^iKO;CDH5+^* and control mice (P21) by whole-mount immunostaining for PECAM1 (green) and LYVE1 (red). **D-F.** Quantification of relative blood vessel density (PECAM1^+^; Control: 1.00 ± 0.01, n = 4; *Vegfr2^iKO;CDH5+^*: 0.90 ± 0.05, n = 4, P = 0.0286), relative lymphatic vessel density (LYVE1^+^; Control: 1.00 ± 0.04, n = 4; *Vegfr2^iKO;CDH5+^*: 0.61 ± 0.05, n = 4, P = 0.0286) and lymphatic vessel diameter (Control: 38.14 ± 1.18 μm, n = 4; *Vegfr2^iKO;CDH5+^*: 30.11 ± 0.71 μm, n = 4, P =0.0286) in B. **G.** Whole-mount immunostaining of ear skins from the *Vegfr2^iKO;CDH5+^*and control mice (P21) for ITGA9 (red) and PROX1. Arrows indicate lymphatic valves. **H-J.** The number of lymphatic endothelial cells was quantified as shown in H and I (PROX1^+^, blind-ended lymphatics, per mm, 134.42 ± 5.94, n = 5; *Vegfr2^iKO;CDH5+^*: 100.25 ± 9.38, n = 5, P = 0.0079; PROX1^+^, collecting lymphatics, per mm, 179.57 ± 7.07, n = 5; *Vegfr2^iKO;CDH5+^*: 122.35 ± 8.12, n = 5, P =0.0079). Lymphatic valves were quantified as shown in J (ITGA9^+^, per grid, Control: 8.85 ± 0.44, n = 5; *Vegfr2^iKO;CDH5+^*: 4.21 ± 0.25, n = 5, P = 0.0079). Scale bar: 200 μm in B; 500 μm in C; 100 μm in G.

### Decrease of lymphatic VEGFR3 expression by the attenuation of VEGFR2

To investigate the mechanism underlying the effects of VEGFR2 insufficiency on lymphatic development, we performed the RNA sequencing analysis of ear skin from the *Vegfr2^iKO;CDH5+^*mutant and control mice (Fig. 2A). The VEGFR2 insufficiency led to a dramatic change in the overall transcriptional profile, including the decreased expression of genes involved in the regulation of lymph/angiogenesis (Fig. 2B-D). The hallmark gene sets, including the hallmarks for lymphangiogenesis and angiogenesis used in the gene set enrichment analysis (GSEA), are mainly defined according to the transcriptome atlas of murine endothelial cells [41]. The top 50 marker genes of lymphangiogenic and angiogenic endothelial cells identified in ten tissues by Kalucka et al [41] were pooled and included in the hallmark gene sets. The hallmark gene sets of other biological processes are based on the molecular signatures database (mh.all.v2024.1.Mm.symbols) [38, 39]. As shown in Fig. 2C, lymphangiogenic genes were significantly downregulated in the skin of *Vegfr2* mutant mice. Heatmaps of the subset of the differentially expressed lymphangiogenic genes are shown in Fig. 2D. There was an obvious decrease in the transcript level of lymphatic regulators including *Vegfr3* and *Prox1* after the endothelial *Vegfr2* deletion (Fig. 2E). Consistently, the VEGFR3 protein was also reduced as shown by the immunostaining analysis of skin at different time points after the induced *Vegfr2* deletion (Fig. 2F and Supplemental Fig. 3). The alteration of VEGFR3 protein was further validated by the western blotting analysis as shown in Fig. 2G-H.

**Fig. 2.**
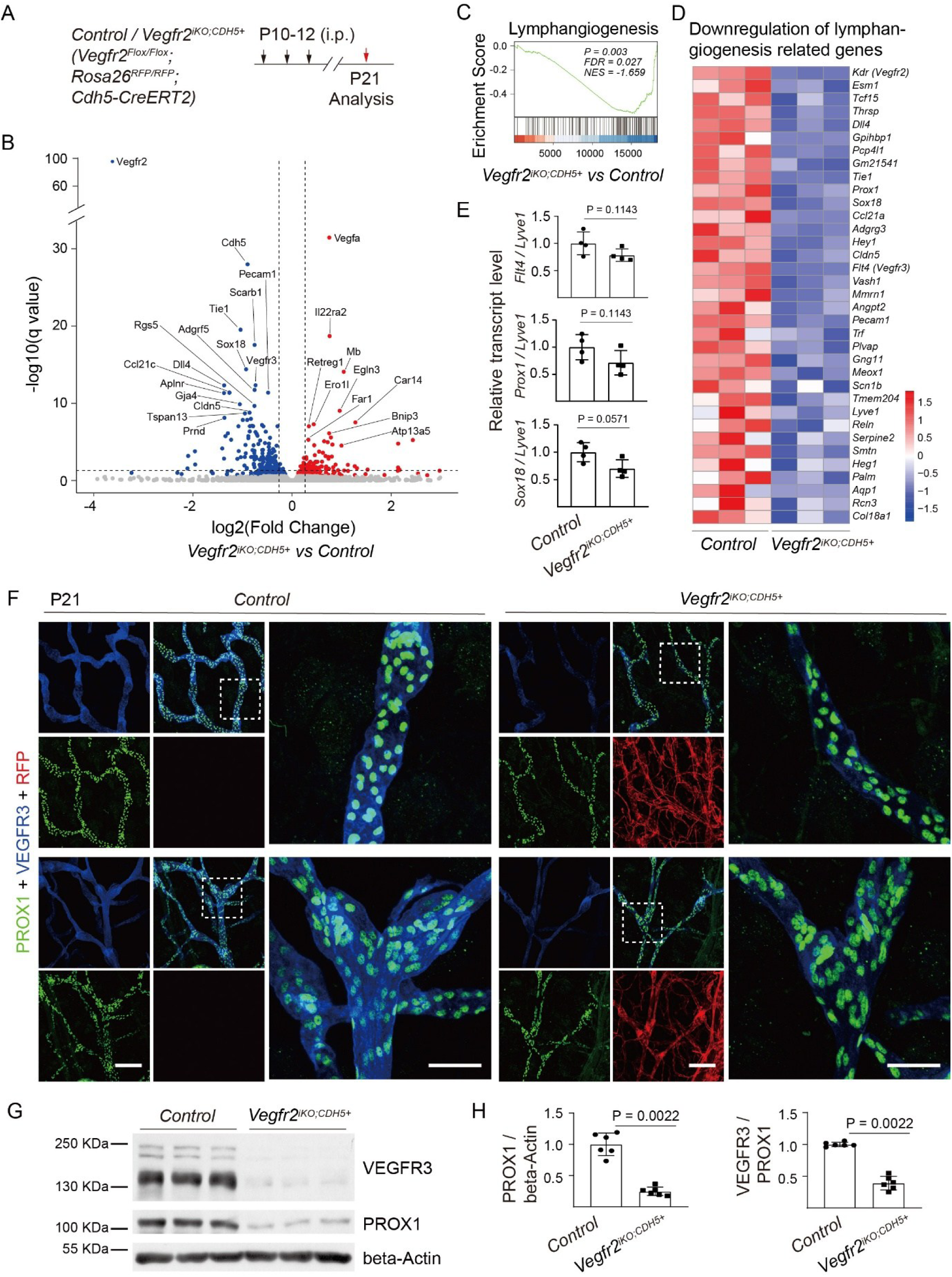
Decrease of the lymphatic VEGFR3 in dermal tissues of *Vegfr2* mutants. **A.** Tamoxifen intraperitoneal administration (i.p.) and mice analysis scheme. **B-D**. RNA-seq analysis of the ear skin tissues of *Vegfr2^iKO;CDH5+^* and control mice (P21). The Volcano plot of the differentially regulated genes was shown in B. GSEA analysis showed the genes involved in lymphangiogenesis were significantly downregulated (C). The subset of the downregulated lymphatic related genes (P < 0.05) was shown as heatmaps in D. **E.** Quantitative expression analysis of *Prox1* and *Vegfr3* in ear skin of *Vegfr2^iKO;CDH5+^* and control mice (P21, *Vegfr3*/*Lyve1*, Control: 1.00 ± 0.21; *Vegfr2^iKO;CDH5+^*: 0.78 ± 0.11, P = 0.1143; *Prox1*/ *Lyve1*, Control: 1.00 ± 0.23; *Vegfr2^iKO;CDH5+^*: 0.71 ± 0.23, P = 0.1143; *Sox18*/*Lyve1*, Control: 1.00 ± 0.17; *Vegfr2^iKO;CDH5+^*: 0.70 ± 0.16, P = 0.0571; n = 4 for each group). **F.** Whole-mount immunostaining of lymphatic vessels in the ear skin from the *Vegfr2^iKO;CDH5+^* and control mice (P21) for PROX1 (green) and VEGFR3 (blue). The reporter RFP signals indicate the Cre (*Cdh5-CreERT2*)-mediated recombination in blood and lymphatic vessels of the *Vegfr2^iKO;CDH5+^*mice. **G-H.** Western blotting analysis and quantification of VEGFR3 and PROX1 protein in ear skin from the *Vegfr2^iKO;CDH5+^* and control mice (P21). (H, PROX1/beta-Actin: Control: 1.00 ± 0.18; *Vegfr2^iKO;CDH5+^*: 0.25 ± 0.07, P = 0.0022; VEGFR3/PROX1: Control: 1.00 ± 0.04; *Vegfr2^iKO;CDH5+^*: 0.36 ± 0.09, P = 0.0022; n = 6 for each group). Scale bar: 50 μm in F.

## Alteration of endothelial transcriptional programs upon the VEGFR2 insufficiency

Consistent with the decrease of lymphatic size and LEC numbers after the induced *Vegfr2* deletion, there were also less button-like junctions in initial lymphatics of skin tissues as shown by the immunostaining analysis for VE-Cadherin (Supplemental Fig. 4A-B). Transcriptional analyses revealed that the expression of lymphatic regulators were downregulated in the skin tissues of *Vegfr2* mutants (Supplemental Fig. 4C). However, pathways related to immune responses were upregulated as shown by the GSEA analysis (Supplemental Fig. 4D). Specifically, these include responses related to interferon-γ/α, inflammatory response and allograft rejection, TNFα via NFκB, IL6-STAT3 and IL2-STAT5 mediated signaling. This was further supported by the GO analysis showing that GO terms enriched for the downregulated vascular genes include biological processes related to the lymph vessel development, sprouting angiogenesis, endothelial cell migration and proliferation, cell-cell and cell-substrate adhesion, and cell junction assembly (Supplemental Fig. 4E). In contrast, pathways related to the regulation of hematopoiesis, endothelial cell apoptotic processes and inflammatory responses were upregulated in the skin tissues of *Vegfr2* mutants (Supplemental Fig. 4F). There was no obvious difference in immune cells detected in the skin by the immunostaining for CD45, or by the flow cytometry analysis of bone marrows or the cell counts in the periphery blood of the *Vegfr2* mutants compared with the controls when analyzed at P21 (Supplemental Fig. 5A-D). This suggests that additional regulators or pathways may be required to switch the EC fate at the postnatal stage.

To further explore the direct endothelial response upon VEGFR2 insufficiency, we performed the phenotype and molecular analysis of skin tissues from the *Vegfr2* mutants and control mice 48 or 96 hours after the induced gene deletion (Fig. 3A; Supplemental Fig. 6A). Interestingly, the decrease of VEGFR3 protein was readily detectable within 96 hours after the induced deletion of endothelial *Vegfr2*, as shown by the immunostaining and the western blotting analysis for VEGFR3 (Supplemental Fig. 6B-E). We also noticed that *Vegfr2* deletion led to a decrease of DLL4 expression in the blood vessels of the skin tissues of mutants, but there was little expression of DLL4 detected in lymphatics at the stage (Supplemental Fig. 6C). The decrease of VEGFR3 protein was further confirmed by the analysis of skin tissues 48 hours after the induced deletion of endothelial *Vegfr2*, as shown in Fig. 3B-D. The altered dermal expression of lymph/angiogenic genes 48 hours after the endothelial deletion of *Vegfr2* (*Vegfr2^iKO;CDH5+^*) was shown by the volcano plot and heatmaps (Fig. 3E-G). Consistent with the findings described above, pathways related to immune responses were upregulated in the *Vegfr2^iKO;CDH5+^* mutants as shown by the GSEA analysis (Fig. 3H), including responses related to interferon-γ/α, inflammatory response and allograft rejection, IL6-JAK_STAT3 and IL2-STAT5 signaling pathways. GO analysis further showed that pathways related to the lymph vessel development was downregulated, while GO terms enriched for the upregulated genes included the biological processes related to hematopoietic progenitor cell differentiation as well as inflammatory responses within 48 hours after the endothelial *Vegfr2* deletion (Fig 3I-J).

**Fig. 3.**
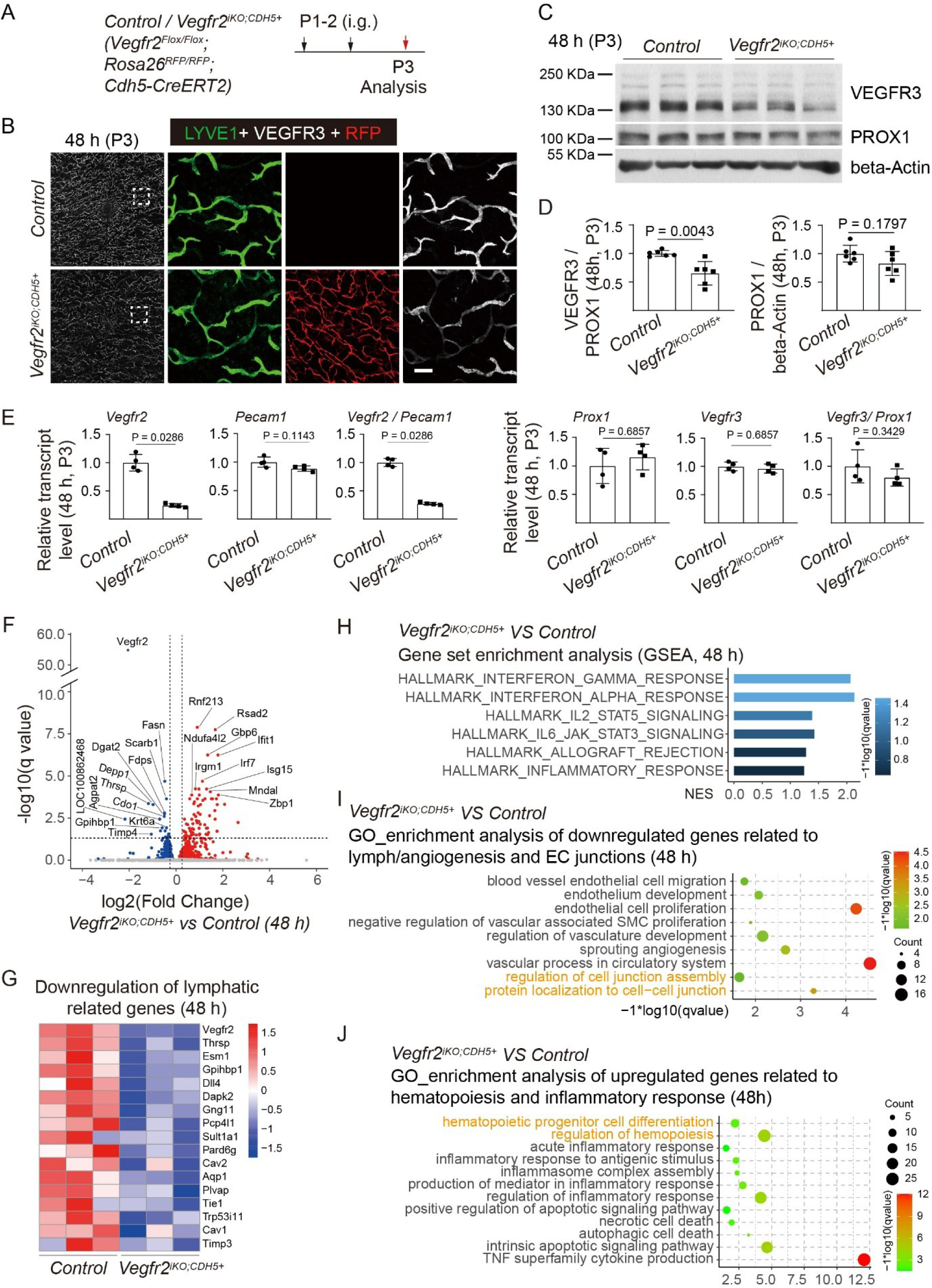
Alteration of lymph/angiogenic gene expression 48 hours after the endothelial *Vegfr2* deletion. **A.** Tamoxifen intragastric administration (i.g.) and mice analysis scheme. **B.** Whole-mount immunostaining of the abdominal skin from the *Vegfr2^iKO;CDH5+^* and control mice (P3) for LYVE1 (green) and VEGFR3 (white). **C-D.** Western blotting analysis and quantification of VEGFR3 and PROX1 protein in back skin from the *Vegfr2^iKO;CDH5+^* and control mice (P3). (D, VEGFR3/PROX1: Control: 1.00 ± 0.05; *Vegfr2^iKO;CDH5+^*: 0.65 ± 0.21, P = 0.0043; PROX1/β-actin: Control: 1.00 ± 0.15; *Vegfr2^iKO;CDH5+^*: 0.83 ± 0.21, P = 0.1797; n = 6 for each group). **E-J.** RNAseq analysis of the abdominal skin tissues of *Vegfr2^iKO;CDH5+^* and control mice (P3). **E.** Quantitative expression analysis of angiogenic and lymphangiogenic genes in the abdominal skin of *Vegfr2^iKO;CDH5+^* and control mice (P3, *Vegfr2*, Control: 1.00 ± 0.15; *Vegfr2^iKO;CDH5+^*: 0.24 ± 0.03, P = 0.0286; *Pecam1*, Control: 1.00 ± 0.09; *Vegfr2^iKO;CDH5+^*: 0.89 ± 0.05, P = 0.1143; *Vegfr2*/*Pecam1*, Control: 1.00 ± 0.07; *Vegfr2^iKO;CDH5+^*: 0.27 ± 0.02, P = 0.0286; *Prox1*, Control: 1.00 ± 0.31; *Vegfr2^iKO;CDH5+^*: 1.15 ± 0.22, P = 0.6857; *Vegfr3*, Control: 1.00 ± 0.08; *Vegfr2^iKO;CDH5+^*: 0.96 ± 0.08, P = 0.6857; *Vegfr3*/*Prox1*, Control: 1.00 ± 0.29; *Vegfr2^iKO;CDH5+^*: 0.80 ± 0.15, P = 0.3429; n = 4 for each group). **F.** The Volcano plot of the differentially regulated genes 48 hours after the endothelial *Vegfr2* deletion. **G.** The subset of the downregulated lymphatic related genes (P < 0.05) was shown as heatmaps. **H**. GSEA analysis showed the increased enrichment in genes related to inflammatory responses when analyzed 48 hours after the endothelial VEGFR2 insufficiency. **I-J.** GO terms enriched for the significantly down-regulated genes were related to vascular development, endothelial cell migration and proliferation, and cell junction assembly (**I**), while significantly upregulated genes were related to hematopoiesis and immune responses when analyzed 48 hours after the induced endothelial deletion of *Vegfr2* (**J**). Scale bar: 100 μm in B.

### Lymphatic inhibition by targeting *Vegfr2* in PROX1^+^ cells

To further characterize the role of lymphatic VEGFR2, we employed an inducible Cre deleter under the control of *Prox1* promoter for the generation of the *Vegfr2* deletion in lymphatics (*Vegfr2^iKO;PROX1+^*). The induced deletion of *Vegfr2* was performed by the tamoxifen treatment at postnatal days 1 to 4 (P1-4) and mice were analyzed at P5-28 (Fig. 4A). The Cre-mediated recombination was confirmed by the expression of reporter RFP in the dermal lymphatic vessels of the *Vegfr2^iKO;CDH5+^* mice (Fig. 4B). Consistent with the phenotypes observed in the ear skin of the *Vegfr2* mutant mice, the lymphatic deletion of *Vegfr2* altered the lymphatic morphology, displaying a decrease of lymphatic vessel size (Fig. 4C-E). VEGFR2 was highly expressed by the lymphatic valves (Fig. 4F) and the loss of lymphatic VEGFR2 also resulted in a significant decrease in the lymphatic valves of pre-collectors (Fig. 4G-H). Consistently, a decrease of VEGFR3 protein was detected in the dermal lymphatic vessels of the *Vegfr2^iKO;PROX1+^* mice when analyzed 96 h after the induced gene deletion (Fig. 4I).

**Fig. 4.**
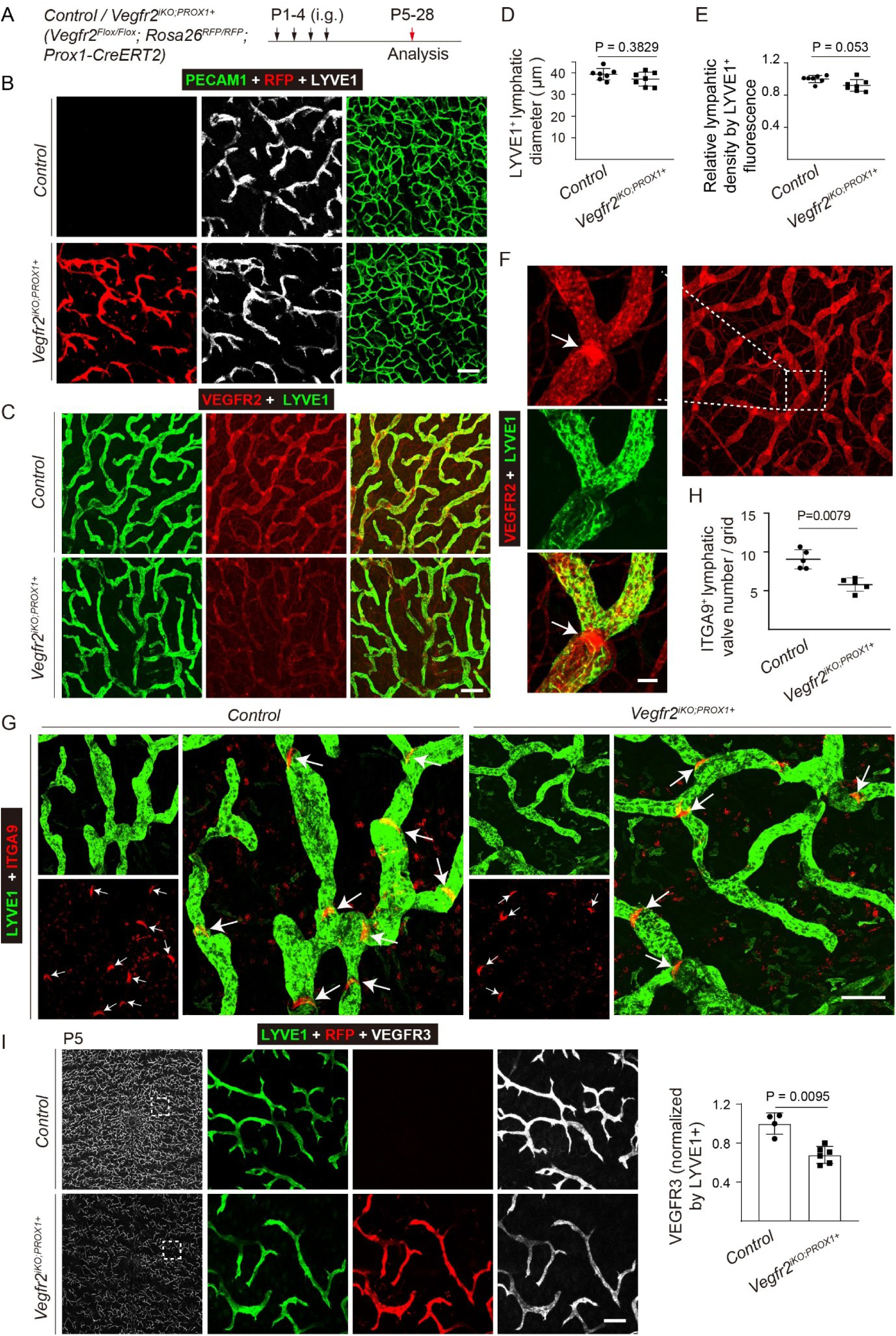
Lymphatic inhibition upon the *Vegfr2* deletion in PROX1^+^ cells. **A.** Tamoxifen intragastric administration (i.g. from P1-4) and mice analysis scheme. **B.** Whole-mount immunostaining of the abdominal skin from the *Vegfr2^iKO;PROX1+^* and control mice (P7) for LYVE1 (grey) and PECAM1 (green). The reporter RFP signals indicate the Cre (*Prox1-CreERT2*)-mediated recombination in lymphatic vessels of the *Vegfr2^iKO;PROX1+^* mice. **C.** Analysis of *Vegfr2* deletion in the dermal lymphatic vessels of the *Vegfr2^iKO;PROX1+^* and control mice (P28) by the whole-mount immunostaining for VEGFR2 (Red) and LYVE1 (green). **D-E.** Quantification of lymphatic diameter was shown in D (Control: 39.34 ± 2.68 μm, n = 7; *Vegfr2^iKO;PROX1+^*: 37.10 ± 3.25 μm, n = 7, P = 0.3829). Relative lymphatic vessel density was shown in E (P21-28, LYVE1^+^; Control: 1.00 ± 0.04, n = 7; *Vegfr2^iKO;PROX1+^*: 0.92 ± 0.07, n = 7, P = 0.0530). **F.** Whole-mount immunostaining of lymphatic vessels (LYVE1, green) in the ear skin to show the high expression of VEGFR2 (red) in the lymphatic valves of pre-collectors (arrows). **G-H.** Analysis of lymphatic valves (arrows) in the ventral side of ear skin from the *Vegfr2^iKO;PROX1+^* and control mice (P28) by whole-mount immunostaining for ITGA9 (Red) and LYVE1 (green). The number of lymphatic valves was quantified as shown in H (ITGA9^+^, per grid, Control: 9.05 ± 1.23, n = 5; *Vegfr2^iKO;PROX1+^*: 5.79 ± 0.87, n = 5, P = 0.0079). **I.** Whole-mount immunostaining of lymphatic vessels in the abdominal skin from the *Vegfr2^iKO;PROX1+^*and control mice (P5) for LYVE1 (green) and VEGFR3 (grey). Quantification of the relative VEGFR3 protein (Control: 1.00 ± 0.11, n = 4; *Vegfr2^iKO;PROX1+^*: 0.68 ± 0.09, n = 6, P = 0.0095). The reporter RFP signals indicate the Cre (*Prox1-CreERT2*)-mediated recombination in lymphatic vessels of the *Vegfr2^iKO;PROX1+^* mice. Scale bar: 100 μm in B, G and I; 200 μm in C; 30 μm in F.

It has been reported that the lymphatic vasculature in ear skins is remodeled from a primary vascular plexus into a mature network composed of blind-ended lymphatic capillaries and valve-containing collecting vessels between P12–P21 [46]. Therefore, the gene deletion was induced from P10 (P10-12 or P10-14), and the *Vegfr2* deletion was shown by the immunostaining analysis for VEGFR2 (Fig. 5A-B). Consistently, the induced lymphatic deletion of *Vegfr2* from P10 also led to a dramatic alteration of the lymphatic morphology including a significant decrease in the lymphatic size and vessel density by the quantification of LYVE1^+^ areas (Fig. 5C-E).

**Fig. 5.**
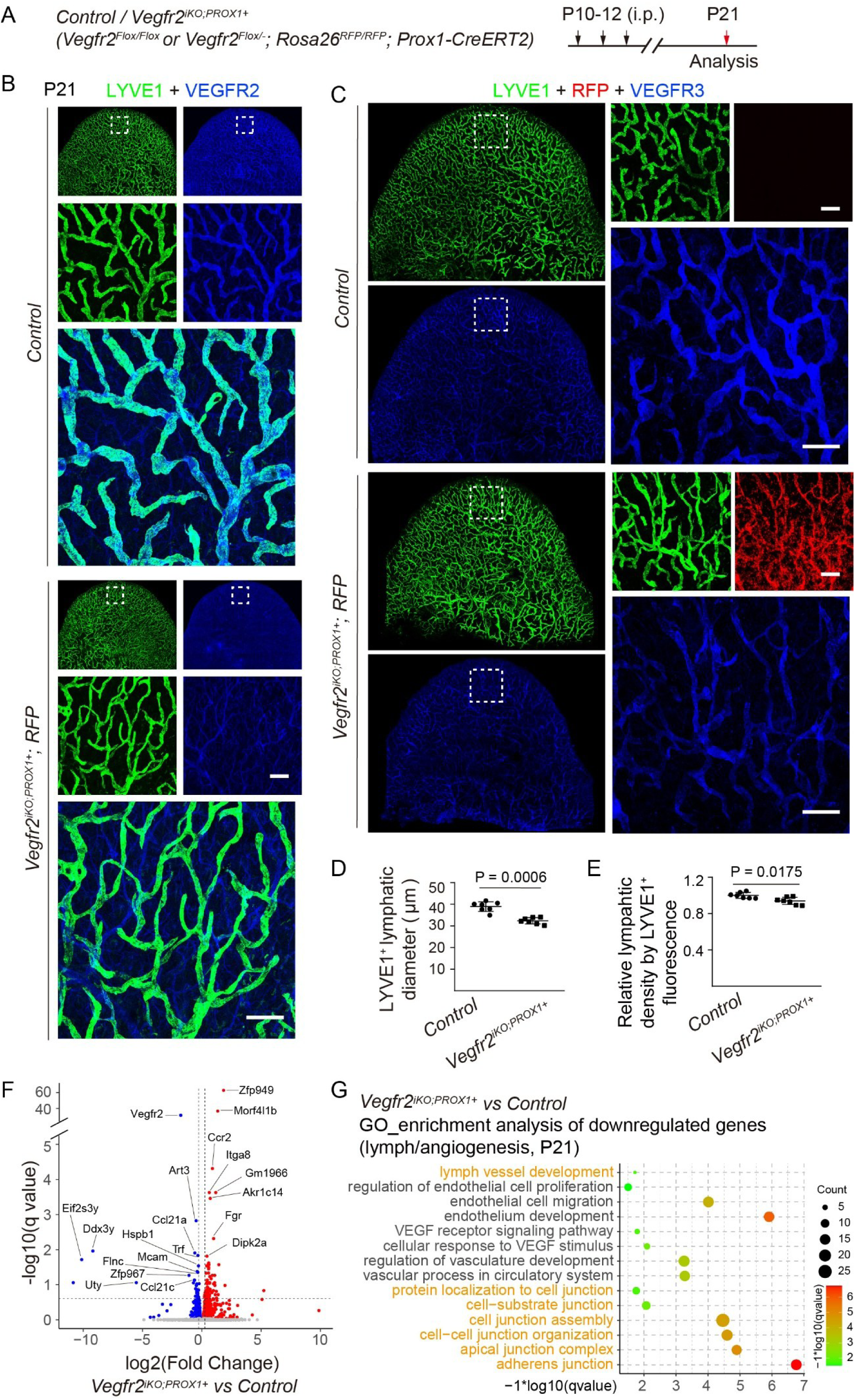
A developmental stage-related effect on lymphatic remodeling by the attenuation of VEGFR2. **A.** Tamoxifen intraperitoneal administration (i.p. from P10-12) and mice analysis scheme. **B.** Analysis of VEGFR2 deletion in the dermal lymphatic vessels of the *Vegfr2^iKO;PROX1+^* and control mice (P21) via whole-mount immunostaining for LYVE1 (green) and VEGFR2 (blue). **C.** Whole-mount immunostaining of lymphatic vessels in the ear skin from the *Vegfr2^iKO;PROX1+^* and control mice (P21) for LYVE1 (green) and VEGFR3 (blue). The reporter RFP signals indicate the Cre (*Prox1-CreERT2*)-mediated recombination in lymphatic vessels of the *Vegfr2^iKO;PROX1+^* mice. **D.** Quantification of lymphatic vessel diameter (Control: 38.92 ± 2.26 μm, n = 7; *Vegfr2^iKO;PROX1+^*: 32.35 ± 1.48 μm, n = 7, P =0.0006). **E.** Quantification of relative lymphatic vessel density (LYVE1^+^, per grid; Control: 1.00 ± 0.03, n = 7; *Vegfr2^iKO;PROX1+^*: 0.94 ± 0.04, n = 7, P = 0.0175). **F.** The volcano plot showed the differentially regulated genes after targeting *Vegfr2* in PROX1^+^ cells. **G.** GO analysis revealed that pathways related to lymph/angiogenesis, particularly endothelial cell migration and proliferation, and cell-cell junction and cell junction assembly were significantly downregulated. Scale bar: 200 μm in B and C.

It is worth pointing out that the lymphatic inhibition in *Vegfr2^iKO;PROX1+^*mice was less severe than that of *Vegfr2^iKO;CDH5+^* mice. It is possible that the endothelial Cre deleter (*Cdh5-CreERT2*) mediated a more efficient gene deletion than that of the lymphatic one (*Prox1-CreERT2*) in LECs. Of note, there was still VEGFR2 remaining in the dermal lymphatics of *Vegfr2^iKO;PROX1+^* mice as evidenced by the immunostaining analysis. On the other hand, the pan-endothelial expression of Cre recombinase under the control of *Cdh5* promoter also mediated an efficient *Vegfr2* deletion in BECs, resulting in the suppression of blood vascular formation. This may indirectly contribute to the lymphatic abnormality of the *Vegfr2^iKO;CDH5+^* mice. Furthermore, we also observed that PROX1 was expressed by a proportion of blood vascular endothelial cells in this study (Supplemental Fig. 7). Using the *Prox1-EGFP* mouse model with the expression of EGFP under the *Prox1* promoter [36], we found that EGFP signals could be detected in a proportion of blood vessels in several tissues examined including head, tail skin, abdominal skin, mesentery and inferior vena cava at the embryonic and postnatal stages (Supplemental Fig. 7A-F). The expression of EGFP in LECs was verified by the immunostaining for PROX1 (Supplemental Fig. 7B, C and F). Further analysis revealed that these EGFP^+^ blood vessels were mainly veins with the negative immunostaining for DLL4 (Supplemental Fig. 7A and E). Consistently, we showed that the Cre deletor-mediated expression of EGFP was detected in a proportion of BECs in veins and capillaries of trachea, intestinal villi and diaphragm using the *Prox1-CreERT2;Rosa26-mTmG* mice (Supplemental Fig. 7G-J). Therefore, apart from the induced deletion of *Vegfr2* in LECs, the deficiency of VEGFR2 could occur in the PROX1^+^ BECs. This may contribute to the alteration of lymphatic morphogenesis resulting from any potential blood vascular effects.

### A transcriptional trend of LEC to hematopoietic transition upon the lymphatic VEGFR2 insufficiency

We performed the RNA-seq analysis of dermal tissues from the *Vegfr2^iKO;PROX1+^* and littermate control mice. The alteration of lymphatic transcriptome was detected by the RNA-seq analysis after the induced deletion of *Vegfr2* in PROX1^+^ cells, showing the decreased expression of genes involved in the EC development and lymphangiogenesis (Fig. 5F-G). Consistent with the transcriptome signature observed in the skin tissues after the pan-endothelial deletion of *Vegfr2*, GSEA analyses identified upregulation of immune pathways in the *Vegfr2^iKO;PROX1+^* mutants, including responses related to interferon-γ/α, inflammatory response and allograft rejection, IL6-JAK_STAT3 and TNFa signaling via NFκB (Fig. 6A-B). This was further supported by the GO analysis showing that GO terms enriched for the upregulated genes include biological processes related to the hematopoietic progenitor cell differentiation, hematopoietic stem cell migration, regulation of hemopoiesis as well as inflammatory response pathways in the skin tissues of *Vegfr2* mutants (Fig. 6C), suggesting the transcriptional trend of LEC to hematopoiesis transition.

**Fig. 6.**
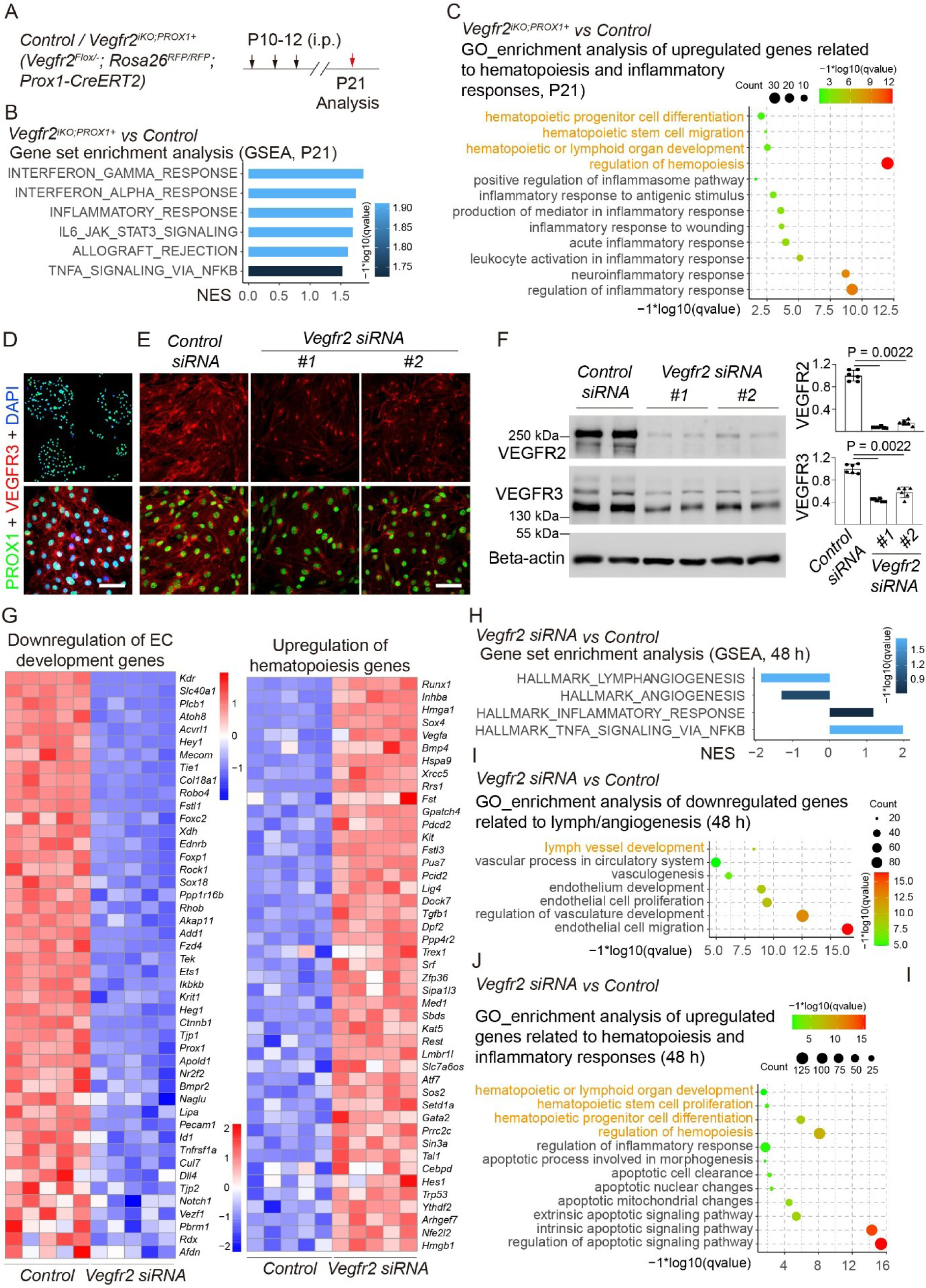
Evidence of endothelial to hematopoietic transition by the lymphatic VEGFR2 insufficiency. **A.** Tamoxifen intraperitoneal administration (i.p.) and analysis scheme. **B**. GSEA analysis showed the increased enrichment in genes related to inflammatory responses after the lymphatic deletion of *Vegfr2*. **C**. GO analysis revealed that pathways related to hematopoiesis and inflammatory responses were significantly upregulated. **D-E.** Immunostaining analysis for PROX1 (green) and VEGFR3 (red) with isolated primary LECs from *tsA58T Ag;Tek-Cre* mice before or after the siRNA-mediated knockdown of *Vegfr2*. **F.** Western blotting analysis of VEGFR2 and VEGFR3 protein in primary LECs after the siRNA-mediated *Vegfr2* knockdown (VEGFR2/beta-Actin: *Control siRNA*: 1.00 ± 0.10; *Vegfr2 siRNA #1*: 0.07 ± 0.01, P = 0.0022; *Vegfr2 siRNA #2*: 0.16 ± 0.05, P = 0.0022; VEGFR3/beta-Actin: *Control siRNA*: 1.00 ± 0.08; *Vegfr2 siRNA #1*: 0.43 ± 0.03, P = 0.0022; *Vegfr2 siRNA #2 siRNA*: 0.57 ± 0.09, P = 0.0022; n = 6 for each group). **G.** Heatmaps showed the subset of the downregulated genes related to EC development and upregulated hematopoiesis related genes (P < 0.01). **H**. GSEA analysis showed the enrichment of downregulated genes related to lymph/angioenesis but upregulated genes to inflammatory responses after the siRNA-mediated knockdown of *Vegfr2*. **I**-**J**. GO analysis revealed that pathways related to lymph/angiogenesis, particularly endothelial cell development were significantly downregulated (**I**), while hematopoiesis, endothelial apoptosis and inflammatory responses related pathways were significantly upregulated (**J**) after the siRNA knockdown of *Vegfr2*. Scale bar: 50 μm in A and B.

To further dissect the functional mechanism of VEGFR2 in lymphatics, we isolated primary lymphatic endothelial cells from the transgenic mice overexpressing tsA58T Ag [44]. The identity of primary LECs was confirmed by the immunostaining analysis for PROX1 and VEGFR3 (Fig. 6D). Using the cultured primary LECs, we demonstrated that there was a significant decrease of VEGFR3 in LECs when VEGFR2 expression was reduced by the siRNA-mediated knockdown (Fig. 6E-F). This suggests a cell autonomous effect on the alteration of VEGFR3 by the lymphatic VEGFR2 insufficiency. Furthermore, we performed the RNA-seq analysis of LECs after the siRNA-mediated knockdown of *Vegfr2*. As shown in Fig. 6G, genes related to the EC development were significantly downregulated while the genes related to the hematopoietic differentiation (e.g. *Runx1*) were upregulated. This was further supported by the GSEA and GO analysis (Fig. 6H-J), showing that the downregulated genes were enriched in the biological processes related to the EC development and lymph/angiogenesis, while the upregulated genes were related to the hematopoiesis and inflammatory response pathways after the siRNA-mediated knockdown of *Vegfr2*.

## Discussion

VEGFR2 mediates a key signaling pathway for the regulation of endothelial cell lineage and angiogenic vessel growth [1, 47, 48]. However, its participation in lymphatic growth and the underlying functional mechanism are not fully characterized. Findings from this study employing the endothelial and lymphatic *Vegfr2* mutant mice as well as the *in vitro* LEC work show that VEGFR2 is required for the lymphatic development and that its insufficiency may alter the LEC identity with a transcriptional trend to become hematopoietic cells.

VEGFR2 is expressed by lymphatic endothelial cells and has been shown to participate in the regulation of developmental as well as pathological lymphangiogenesis. It was reported that the conditional deletion of *Vegfr2* in endothelial cells using a *Lyve1-Cre* transgenic mouse line led to the lymphatic hypoplasia in skin tisses [27]. As LYVE1 is also expressed in some blood vascular endothelial cells (for example in yolk sac BECs) [20, 49], the disruption of embryonic angiogenesis after the loss of endothelial VEGFR2 may contribute to the underdeveloped dermal lymphatic network. Furthermore, mice carrying a mutation in the SEMA3A binding site of NRP1, a coreceptor of VEGFR2, were shown to display lymphatic valve defects [50, 51]. Enhanced VEGFR2 signaling by the loss of NRP1 or FLT1 receptors was shown to induce lacteal junction zippering [52]. In pathological conditions, VEGFA-VEGFR2 signaling was shown to induce lymphatic growth in cutaneous leishmaniasis [53]. In a mouse corneal suturing model, the expression of soluble VEGFR2 (sKDR) by the splice-shifting morpholinos or the systemic administration of neutralizing antibodies against VEGFR2 suppressed corneal angiogenesis and lymphangiogenesis [54, 55]. VEGFA and the specific VEGFR3 activator VEGFC156S were shown to induce a similar set of target genes as well as some specific ones in primary human lymphatic endothelial cells [56]. Overexpression of VEGFA or VEGFE via an adenoviral vector induced abnormal lymphangiogenesis [28, 29]. It was also found that VEGFA could induce lymphangiogenesis by recruiting bone marrow-derived macrophages to secrete lymphangiogenic factors [57–59] or by the upregulation of VEGFC in vascular endothelial cells [60]. In addition, VEGFC has been shown to promote the VEGFR2/VEGFR3 heterodimerization, which may mediate signals for lymphangiogenesis in cauterised cornea and tumor [22, 61]. It was also reported that VEGFR2 polymorphism (rs2071559, T/C) had an association with lymphatic metastasis in patients with nasopharyngeal carcinoma [62]. Tyrosine kinase inhibitors or neutralizing antibodies against VEGFR2 were reported to have a suppressive effect on tumor-associated lymphatics [63]. However, over-activation of VEGFR2 by excess VEGF ligands or suppression of VEGFR2-mediated signaling could affect blood vascular growth and produce indirect effects on the lymphatic system.

To further investigate specific roles of VEGFR2 in lymphangiogenesis, we employed in this study a mouse strain with the induced deletion of *Vegfr2* in PROX1^+^ cells (*Vegfr2^iKO-PROX1+^*) [32]. PROX1 was expressed mainly in lymphatic endothelial cells within the vascular system [64]. We show that the lymphatic VEGFR2 insufficiency resulted in the suppression of dermal lymphatic vessel growth as well as valve formation in pre-collectors, suggesting a direct role of VEGFR2 in lymphatics. Notably, as observed in this study, PROX1 also marks a proportion of blood vascular endothelial cells in veins and capillaries during the embryogenic and postnatal development. Therefore, apart from the induced deletion of *Vegfr2* in LECs, the induced deletion of *Vegfr2* could occur in the PROX1^+^ BECs, potentially affecting the blood vascular formation or function. In addition, we observed a more potent suppression of dermal lymphatic growth in the *Vegfr2^iKO;CDH5+^* mutants than that of *Vegfr2^iKO;PROX1+^* mice. Therefore, it is possible that the VEGFR2 insufficiency in specific vessel types or regions may also contribute to the defective lymphatic vessel formation.

Related to the lymphatic phenotypes upon the VEGFR2 insufficiency, it is worth emphasizing that the lymphatic VEGFR3 is dramatically reduced in mutant mice with the induced *Vegfr2* deletion in ECs (CDH5^+^) or mainly LECs (PROX1^+^). Consistently, the decrease of VEGFR3 after the endothelial deletion of *Vegfr2* was also observed in blood vascular endothelial cells [65–68]. The alteration of lymphatic VEGFR3 was also reported in mutant mice with the attenuation of ANGPT2-TIE receptor-mediated signaling pathway [69]. In line with this, in a LPS-induced sepsis model, VEGFR3 positive vessel areas and lymphatic diameters were significantly decreased in the lung and draining lymph nodes within 48 hours after the treatment [70]. It was also reported that the Alzheimer’s disease-linked protease BACE2 cleaves the full-length VEGFR3 to attenuate its signaling [71]. Evidences from the above studies point to the fact that the endothelial or lymphatic expression of VEGFR3 could be modulated by various factors in physiological or pathological conditions.

One key question is that how the lymphatic VEGFR2 insufficiency leads to the alteration of VEGFR3 expression in dermal lymphatic vessels as observed in this study. VEGFR2-mediated signaling pathway is a key determinant of EC differentiation and functions [1]. It is therefore possible that the insufficient endothelial VEGFR2-mediated signals may alter the LEC identity as shown by the loss of lymphatic surface marker VEGFR3, which mediates a key signaling pathway for the lymphatic growth, LEC junction formation and valve morphogenesis. It has been shown recently that embryonic lymphatic ECs possess the haemogenic capacity, which is suppressed by the LEC fate determining factor PROX1 [33, 72]. One may wonder whether the insufficient VEGFR2 signaling may also lead to the endothelial-to-hematopoietic transition. Indeed, we observed in this study that the pan-endothelial or lymphatic VEGFR2 insufficiency altered the transcriptional programs, with the increased expression of hematopoietic genes observed in the skin tissues of *Vegfr2* mutant mice while genes involved in the EC development was downregulated by the RNA-seq analysis. Consistently, using the cultured primary LECs, we validated the decrease of VEGFR3 in LECs when VEGFR2 expression was reduced by the siRNA-mediated knockdown. The RNA-seq analysis of LECs after the siRNA-mediated knockdown of *Vegfr2* demonstrated that genes related to the EC development were significantly downregulated while those related to the hematopoietic differentiation (e.g. *Runx1*) were upregulated. Findings from this study imply that VEGFR2 is required for the maintenance of endothelial identity and that its insufficiency may lead to a transcriptional trend of LEC to hematopoietic transition to diminish the VEGFR3-mediated lymphangiogenesis (Supplemental Fig. 8).

Furthermore, we also observed that GO terms enriched for the upregulated dermal genes include biological processes related to the inflammatory responses in the *Vegfr2* mutants or in the LECs with the siRNA-mediated knockdown of *Vegfr2* by the RNA-seq analysis. This may reflect the alteration of the LEC identity. Activation of innate immunity was shown to induce a state of epigenetic plasticity in which external cues can guide the cell transdifferentiation [47]. Studies in this direction will provide further insights into the molecular cues underlying the lymphatic endothelial cell fate transition.

In summary, we show in this study that loss of VEGFR2 in the ECs (CDH5^+^) or LECs (PROX1^+^) may trigger the alteration of the LEC identity as shown by the transcriptional programs, particularly the decreased lymphatic VEGFR3, leading to the defective lymphatic morphogenesis. This implies that targeting endothelial VEGFR2 could affect both angiogenesis and lymphangiogenesis in development and diseases. On the one hand, VEGFR2 could be employed for modulating both blood vascular and lymphatic growth in pathological conditions including cancer, particularly with a therapeutic potential for blocking the tumor cell dissemination via lymphatic as well as blood vascular routes. On the other hand, blockade of VEGFR2 mediated signals could affect the lymphatic transcriptional programs, potentially harmful for the lymphatic growth and the maintenance of preexisting lymphatic networks in physiological conditions.

## Acknowledgement

We thank Dr. Janet Rossant for kindly providing us the *Vegfr2^Flox/Flox^* mouse line, Dr. Yoshiaki Kubota of Keio University for the *Cdh5-CreERT2* mouse line, Dr. Taija Makinen of Uppsala University & Helsinki University for the *Prox1-CreERT2* mouse line, Dr. Bin Zhou of University of Chinese Academy of Sciences for the *Rosa26^RFP^* mouse line, and staff in the Animal facility of Soochow University for technical assistance.

## Reference

1. Lv, J., et al., Epigenetic landscape reveals MECOM as an endothelial lineage regulator. Nat Commun, 2023. 14(1): p. 2390.

2. Lee, S., et al., Prox1 physically and functionally interacts with COUP-TFII to specify lymphatic endothelial cell fate. Blood, 2009. 113(8): p. 1856–9.

3. Wigle, J.T., et al., An essential role for Prox1 in the induction of the lymphatic endothelial cell phenotype. EMBO J, 2002. 21(7): p. 1505–13.

4. Hong, Y.K., et al., Prox1 is a master control gene in the program specifying lymphatic endothelial cell fate. Dev Dyn, 2002. 225(3): p. 351–7.

5. Srinivasan, R.S., et al., The nuclear hormone receptor Coup-TFII is required for the initiation and early maintenance of Prox1 expression in lymphatic endothelial cells. Genes Dev, 2010. 24(7): p. 696–707.

6. Ma, W., et al., Mitochondrial respiration controls the Prox1-Vegfr3 feedback loop during lymphatic endothelial cell fate specification and maintenance. Sci Adv, 2021. 7(18).

7. Alitalo, K., T. Tammela, and T.V. Petrova, Lymphangiogenesis in development and human disease. Nature, 2005. 438(7070): p. 946–53.

8. Petrova, T.V. and G.Y. Koh, Biological functions of lymphatic vessels. Science, 2020. 369(6500).

9. Oliver, G. and M. Detmar, The rediscovery of the lymphatic system: old and new insights into the development and biological function of the lymphatic vasculature. Genes & Development, 2002. 16(7): p. 773–783.

10. Potente, M. and T. Makinen, Vascular heterogeneity and specialization in development and disease. Nat Rev Mol Cell Biol, 2017. 18(8): p. 477–494.

11. Alitalo, K., The lymphatic vasculature in disease. Nat Med, 2011. 17(11): p. 1371–80.

12. Koch, S., et al., Signal transduction by vascular endothelial growth factor receptors. Biochem J, 2011. 437(2): p. 169–83.

13. Jeltsch, M., et al., Receptor tyrosine kinase-mediated angiogenesis. Cold Spring Harb Perspect Biol, 2013. 5(9).

14. Adams, R.H. and K. Alitalo, Molecular regulation of angiogenesis and lymphangiogenesis. Nat Rev Mol Cell Biol, 2007. 8(6): p. 464–78.

15. Ferrara, N., H.P. Gerber, and J. LeCouter, The biology of VEGF and its receptors. Nat Med, 2003. 9(6): p. 669–76.

16. Secker, G.A. and N.L. Harvey, Regulation of VEGFR Signalling in Lymphatic Vascular Development and Disease: An Update. Int J Mol Sci, 2021. 22(14).

17. Karaman, S., V.M. Leppanen, and K. Alitalo, Vascular endothelial growth factor signaling in development and disease. Development, 2018. 145(14).

18. Makinen, T., et al., Inhibition of lymphangiogenesis with resulting lymphedema in transgenic mice expressing soluble VEGF receptor-3. Nat Med, 2001. 7(2): p. 199–205.

19. Karkkainen, M.J., et al., Vascular endothelial growth factor C is required for sprouting of the first lymphatic vessels from embryonic veins. Nat Immunol, 2004. 5(1): p. 74–80.

20. Zhang, L., et al., VEGFR-3 ligand-binding and kinase activity are required for lymphangiogenesis but not for angiogenesis. Cell Res, 2010. 20(12): p. 1319–31.

21. Zhang, Y., et al., Heterogeneity in VEGFR3 levels drives lymphatic vessel hyperplasia through cell-autonomous and non-cell-autonomous mechanisms. Nat Commun, 2018. 9(1): p. 1296.

22. Dixelius, J., et al., Ligand-induced vascular endothelial growth factor receptor-3 (VEGFR-3) heterodimerization with VEGFR-2 in primary lymphatic endothelial cells regulates tyrosine phosphorylation sites. J Biol Chem, 2003. 278(42): p. 40973–9.

23. Deng, Y., X. Zhang, and M. Simons, Molecular controls of lymphatic VEGFR3 signaling. Arterioscler Thromb Vasc Biol, 2015. 35(2): p. 421–9.

24. Nilsson, I., et al., VEGF receptor 2/-3 heterodimers detected in situ by proximity ligation on angiogenic sprouts. Embo J, 2010. 29(8): p. 1377–88.

25. Albuquerque, R.J., et al., Alternatively spliced vascular endothelial growth factor receptor-2 is an essential endogenous inhibitor of lymphatic vessel growth. Nat Med, 2009. 15(9): p. 1023–30.

26. Vogrin, A.J., et al., Evolutionary Differences in the Vegf/Vegfr Code Reveal Organotypic Roles for the Endothelial Cell Receptor Kdr in Developmental Lymphangiogenesis. Cell Rep, 2019. 28(8): p. 2023–2036 e4.

27. Dellinger, M.T., et al., Vascular endothelial growth factor receptor-2 promotes the development of the lymphatic vasculature. PLoS One, 2013. 8(9): p. e74686.

28. Nagy, J.A., et al., Vascular permeability factor/vascular endothelial growth factor induces lymphangiogenesis as well as angiogenesis. J Exp Med, 2002. 196(11): p. 1497–506.

29. Wirzenius, M., et al., Distinct vascular endothelial growth factor signals for lymphatic vessel enlargement and sprouting. J Exp Med, 2007. 204(6): p. 1431–40.

30. Li, X., et al., VEGFR2 pY949 signalling regulates adherens junction integrity and metastatic spread. Nat Commun, 2016. 7: p. 11017.

31. Okabe, K., et al., Neurons Limit Angiogenesis by Titrating VEGF in Retina. Cell, 2014. 159(3): p. 584–596.

32. Bazigou, E., et al., Genes regulating lymphangiogenesis control venous valve formation and maintenance in mice. J Clin Invest, 2011. 121(8): p. 2984–92.

33. Kazenwadel, J., et al., A Prox1 enhancer represses haematopoiesis in the lymphatic vasculature. Nature, 2023. 614(7947): p. 343–348.

34. Haigh, J.J., et al., Cortical and retinal defects caused by dosage-dependent reductions in VEGF-A paracrine signaling. Dev Biol, 2003. 262(2): p. 225–41.

35. Liu, Q., et al., Genetic targeting of sprouting angiogenesis using Apln-CreER. Nat Commun, 2015. 6: p. 6020.

36. Choi, I., et al., Visualization of lymphatic vessels by Prox1-promoter directed GFP reporter in a bacterial artificial chromosome-based transgenic mouse. Blood, 2011. 117(1): p. 362–5.

37. Cao, X., et al., Endothelial TIE1 Restricts Angiogenic Sprouting to Coordinate Vein Assembly in Synergy With Its Homologue TIE2. Arterioscler Thromb Vasc Biol, 2023. 43(8): p. e323–e338.

38. Subramanian, A., et al., Gene set enrichment analysis: a knowledge-based approach for interpreting genome-wide expression profiles. Proc Natl Acad Sci U S A, 2005. 102(43): p. 15545–50.

39. Liberzon, A., et al., The Molecular Signatures Database (MSigDB) hallmark gene set collection. Cell Syst, 2015. 1(6): p. 417–425.

40. Morgan M, F.S., Gentleman R, GSEABase: Gene set enrichment data structures and methods. Bioconductor R package version 1.56.0. 10.18129/B9.bioc.GSEABase. 2021.

41. Kalucka, J., et al., Single-Cell Transcriptome Atlas of Murine Endothelial Cells. Cell, 2020. 180(4): p. 764–779 e20.

42. Shen, B., et al., Genetic dissection of tie pathway in mouse lymphatic maturation and valve development. Arterioscler Thromb Vasc Biol, 2014. 34(6): p. 1221–30.

43. Kazenwadel, J., M.Z. Michael, and N.L. Harvey, Prox1 expression is negatively regulated by miR-181 in endothelial cells. Blood, 2010. 116(13): p. 2395–401.

44. Yamaguchi, T., et al., Development of a new method for isolation and long-term culture of organ-specific blood vascular and lymphatic endothelial cells of the mouse. FEBS J, 2008. 275(9): p. 1988–98.

45. Koni, P.A., et al., Conditional vascular cell adhesion molecule 1 deletion in mice: impaired lymphocyte migration to bone marrow. J Exp Med, 2001. 193(6): p. 741–54.

46. Lutter, S., et al., Smooth muscle-endothelial cell communication activates Reelin signaling and regulates lymphatic vessel formation. J Cell Biol, 2012. 197(6): p. 837–49.

47. Sayed, N., et al., Transdifferentiation of human fibroblasts to endothelial cells: role of innate immunity. Circulation, 2015. 131(3): p. 300–9.

48. Taura, D., et al., Induction and isolation of vascular cells from human induced pluripotent stem cells--brief report. Arterioscler Thromb Vasc Biol, 2009. 29(7): p. 1100–3.

49. Mouta Carreira, C., et al., LYVE-1 is not restricted to the lymph vessels: expression in normal liver blood sinusoids and down-regulation in human liver cancer and cirrhosis. Cancer Res, 2001. 61(22): p. 8079–84.

50. Bouvree, K., et al., Semaphorin3A, Neuropilin-1, and PlexinA1 are required for lymphatic valve formation. Circ Res, 2012. 111(4): p. 437–45.

51. Jurisic, G., et al., An unexpected role of semaphorin3a-neuropilin-1 signaling in lymphatic vessel maturation and valve formation. Circ Res, 2012. 111(4): p. 426–36.

52. Zhang, F., et al., Lacteal junction zippering protects against diet-induced obesity. Science, 2018. 361(6402): p. 599–603.

53. Weinkopff, T., et al., Leishmania major Infection-Induced VEGF-A/VEGFR-2 Signaling Promotes Lymphangiogenesis That Controls Disease. J Immunol, 2016. 197(5): p. 1823–31.

54. Uehara, H., et al., Dual suppression of hemangiogenesis and lymphangiogenesis by splice-shifting morpholinos targeting vascular endothelial growth factor receptor 2 (KDR). FASEB J, 2013. 27(1): p. 76–85.

55. Yuen, D., B. Pytowski, and L. Chen, Combined blockade of VEGFR-2 and VEGFR-3 inhibits inflammatory lymphangiogenesis in early and middle stages. Invest Ophthalmol Vis Sci, 2011. 52(5): p. 2593–7.

56. Dieterich, L.C., et al., Distinct transcriptional responses of lymphatic endothelial cells to VEGFR-3 and VEGFR-2 stimulation. Sci Data, 2017. 4: p. 170106.

57. Cursiefen, C., et al., VEGF-A stimulates lymphangiogenesis and hemangiogenesis in inflammatory neovascularization via macrophage recruitment. J Clin Invest, 2004. 113(7): p. 1040–50.

58. Maruyama, K., et al., Inflammation-induced lymphangiogenesis in the cornea arises from CD11b-positive macrophages. J Clin Invest, 2005. 115(9): p. 2363–72.

59. Schoppmann, S.F., et al., Tumor-associated macrophages express lymphatic endothelial growth factors and are related to peritumoral lymphangiogenesis. Am J Pathol, 2002. 161(3): p. 947–56.

60. Nakao, S., et al., Blood vessel endothelial VEGFR-2 delays lymphangiogenesis: an endogenous trapping mechanism links lymph- and angiogenesis. Blood, 2011. 117(3): p. 1081–90.

61. Durre, T., et al., uPARAP/Endo180 receptor is a gatekeeper of VEGFR-2/VEGFR-3 heterodimerisation during pathological lymphangiogenesis. Nat Commun, 2018. 9(1): p. 5178.

62. Hu, K., et al., Association of the rs2071559 (T/C) polymorphism with lymphatic metastasis in patients with nasopharyngeal carcinoma. Oncol Lett, 2017. 14(6): p. 7681–7686.

63. Padera, T.P., et al., Differential response of primary tumor versus lymphatic metastasis to VEGFR-2 and VEGFR-3 kinase inhibitors cediranib and vandetanib. Mol Cancer Ther, 2008. 7(8): p. 2272–9.

64. Wigle, J.T. and G. Oliver, Prox1 function is required for the development of the murine lymphatic system. Cell, 1999. 98(6): p. 769–78.

65. Benedito, R., et al., Notch-dependent VEGFR3 upregulation allows angiogenesis without VEGF–VEGFR2 signalling. Nature, 2012. 484(7392): p. 110–114.

66. Jang, J.Y., et al., VEGFR2 but not VEGFR3 governs integrity and remodeling of thyroid angiofollicular unit in normal state and during goitrogenesis. EMBO Molecular Medicine, 2017. 9(6): p. 750–769.

67. Zarkada, G., et al., VEGFR3 does not sustain retinal angiogenesis without VEGFR2. Proceedings of the National Academy of Sciences, 2015. 112(3): p. 761–766.

68. Tammela, T., et al., Blocking VEGFR-3 suppresses angiogenic sprouting and vascular network formation. Nature, 2008. 454(7204): p. 656–660.

69. Korhonen, E.A., et al., Lymphangiogenesis requires Ang2/Tie/PI3K signaling for VEGFR3 cell-surface expression. J Clin Invest, 2022. 132(15).

70. Zhang, P.H., et al., Efficient pulmonary lymphatic drainage is necessary for inflammation resolution in ARDS. JCI Insight, 2024. 9(1).

71. Schmidt, A., et al., The Alzheimer’s disease-linked protease BACE2 cleaves VEGFR3 and modulates its signaling. J Clin Invest, 2024.

72. Mäkinen, T., Hemogenic activity of lymphatic endothelium unleashed. Nature Cardiovascular Research, 2023. 2(3): p. 230–231.

