## Supplemental Fig. 1-8 for "Lymphatics unload VEGFR3 in response to the alteration of VEGFR2-mediated transcriptional programs"

Running title: Requirement of VEGFR2 for lymphatic endothelial identity

* Correspondence should be addressed to Dr. Yulong He

Key words: VEGFR2, VEGFR3, transcriptional programs, endothelial cell lineage, lymphatic endothelial cell, lymphangiogenesis

**Supplemental Figures**

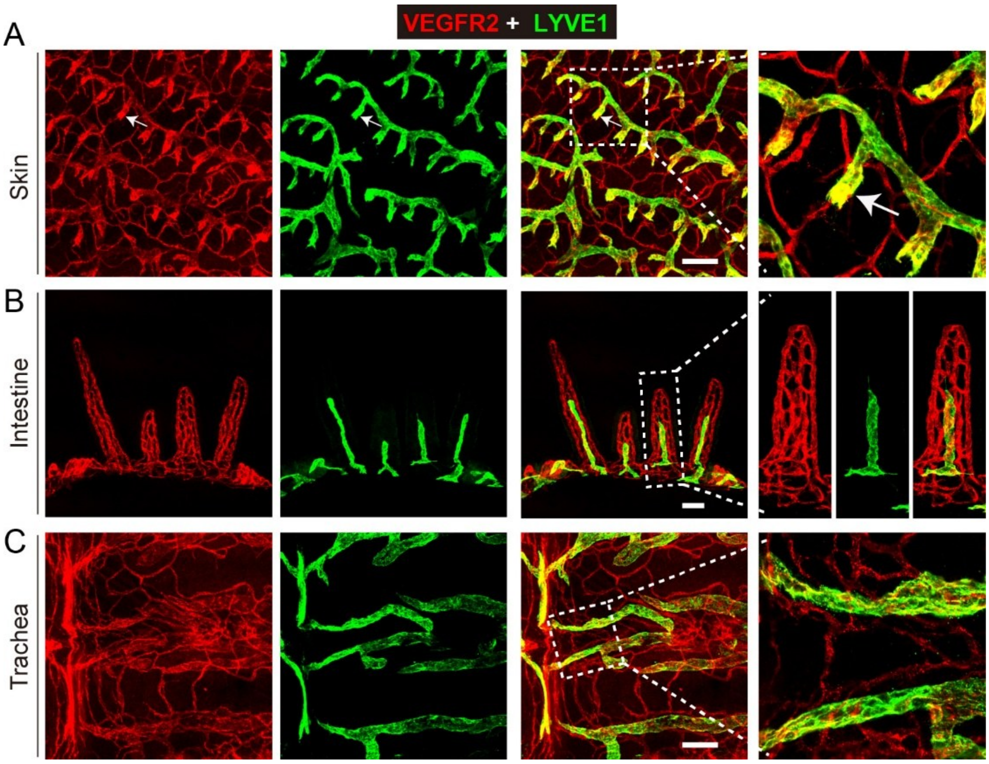


**Supplemental Fig. 1 Analysis of VEGFR2 expression in lymphatics of different tissues. A-C**. Whole-mount immunostaining of VEGFR2 expression in lymphatic vessels of the ventral skin (A), villi (B) and trachea (C) in wild-type mice (P7; VEGFR2, red; LYVE1, green). Arrows indicate the high expression of VEGFR2 in dermal initial lymphatic vessels. Scale bar: 100 μm in A and C; 200 μm in B.

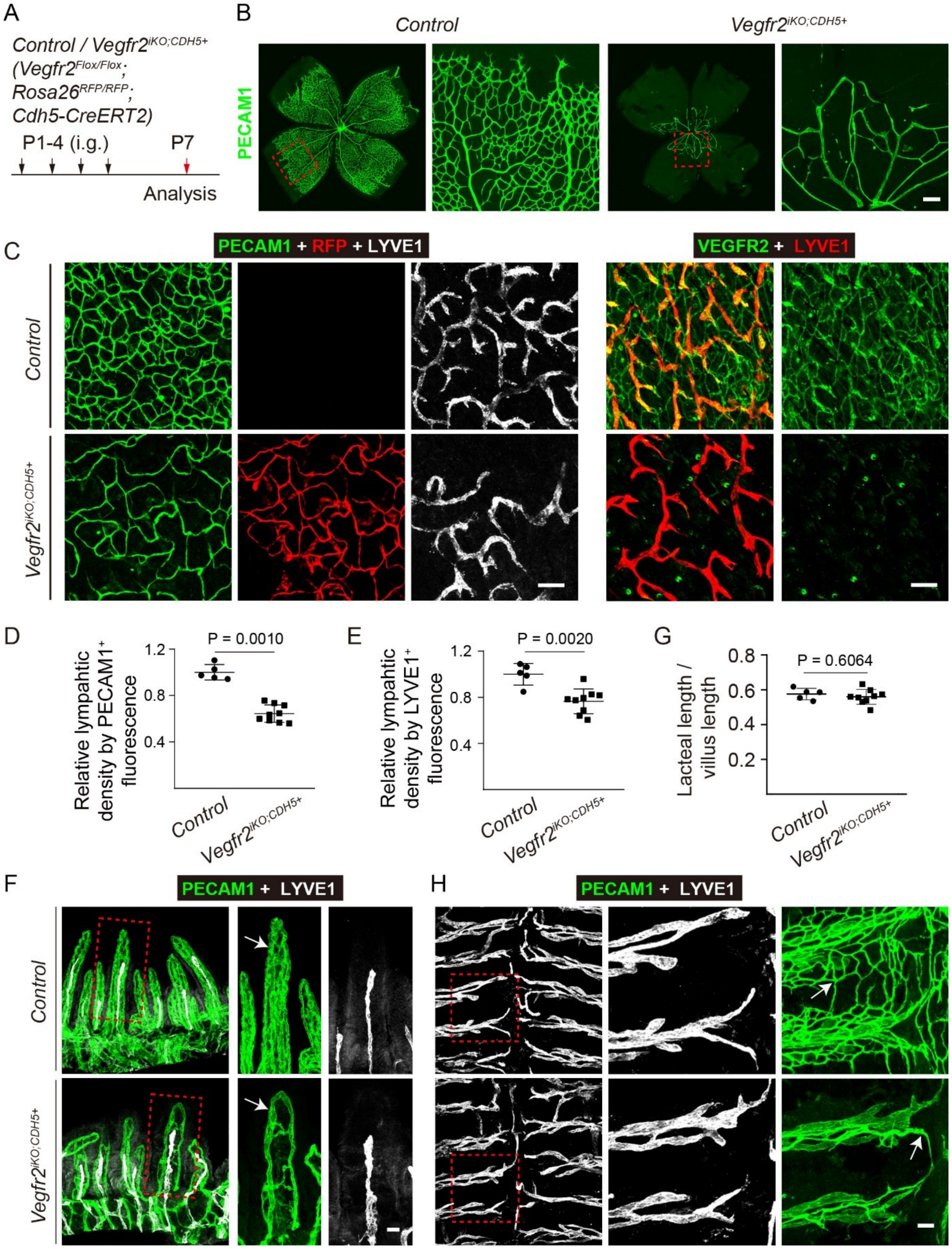


**Supplemental Fig. 2 Suppression of both angiogenesis and lymphangiogenesis after the postnatal pan-endothelial deletion of *Vegfr2*. A**. Tamoxifen intragastric administration (i.g.) and mice analysis scheme. **B**. Analysis of blood vessels in retina from the *Vegfr2*$^{iKO;CDH5+}$ and control mice (P7) by whole-mount immunostaining for PECAM1 (green). **C.** Analysis of blood vessels and lymphatic vessels in the abdominal skin from the *Vegfr2*$^{iKO;CDH5+}$ and control mice (P7) by whole-mount immunostaining for PECAM1 (green) and LYVE1 (grey, left panel), and for VEGFR2 (green) and LYVE1 (red, right panel). The reporter RFP signals indicate the expression of reporter gene (tdTomato) by the Cre (*Cdh5-CreERT2*)-mediated recombination in blood and lymphatic vessels of the *Vegfr2*$^{iKO;CDH5+}$ mice. **D-E**. Quantification of relative blood vessel area (D, PECAM1$^{+}$, Control: 1.00 ± 0.07, n = 5; *Vegfr2*$^{iKO;CDH5+}$: 0.64 ± 0.07, n = 9, P = 0.0010) and lymphatic vessel area (E, LYVE1$^{+}$, Control: 1.00 ± 0.09, n = 5; *Vegfr2*$^{iKO;CDH5+}$: 1.00 ± 0.11, n = 9, P = 0.0020). **F-H.** Analysis of blood vessels and lymphatic vessels in the intestinal villi (F) and trachea (H) from the *Vegfr2*$^{iKO;CDH5+}$ and control mice (P7) by whole-mount immunostaining for PECAM1 (green) and LYVE1 (grey). Quantification of the ratio of lacteal to villus length was shown in G (Control: 0.58 ± 0.03, n = 5; *Vegfr2*$^{iKO;CDH5+}$: 0.56 ± 0.04, n = 9, P = 0.6064). Scale bar: 100 μm in B, C; 30 μm in F; 50 μm in H.

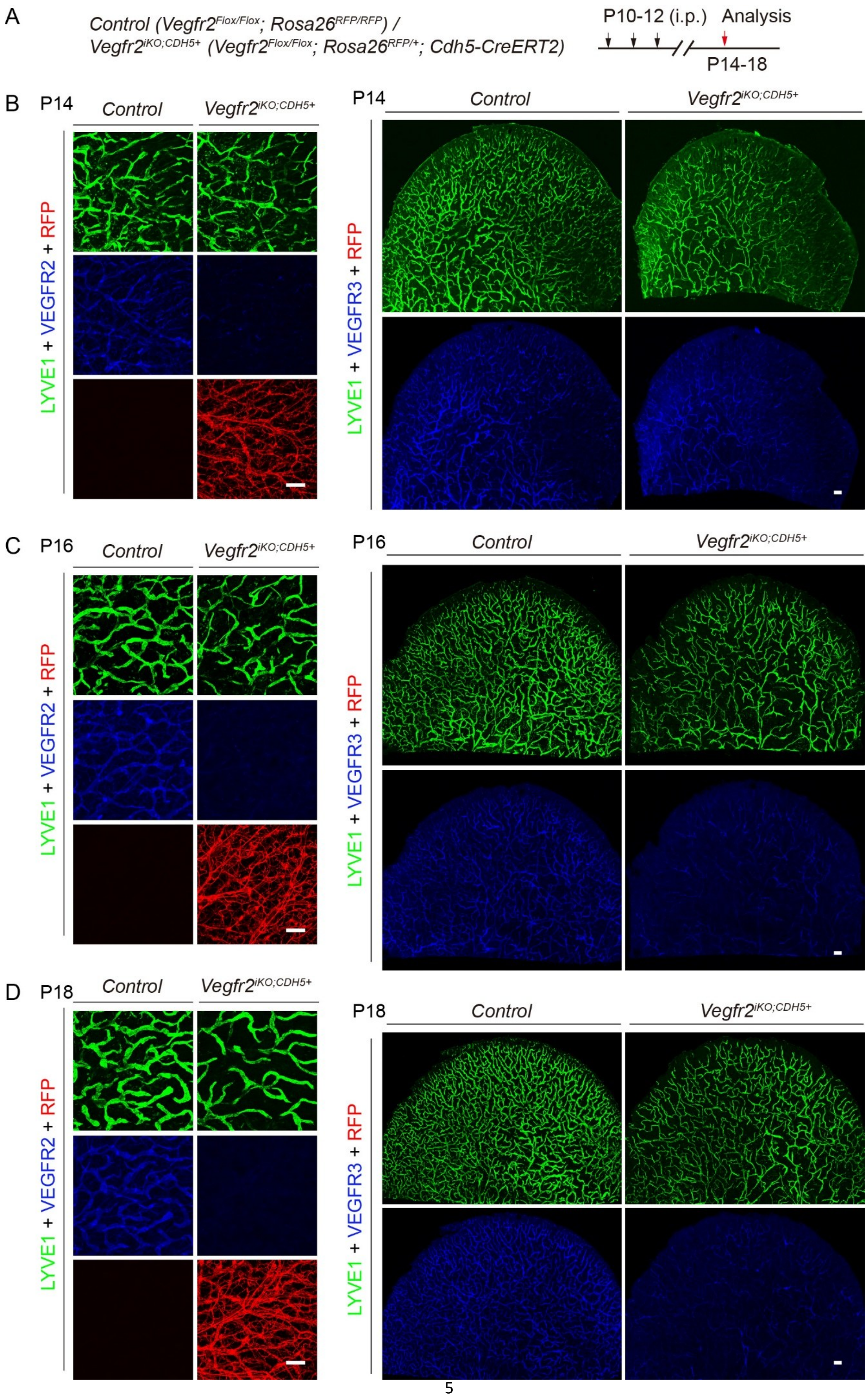

A
Control (Vegfr2Flox/Flox; Rosa26RFP/RFP) /
Vegfr2iKO;CDH5+ (Vegfr2Flox/Flox; Rosa26RFP/+; Cdh5-CreERT2)
P10-12 (i.p.)
Analysis
P14-18
B
P14
Control
Vegfr2iKO;CDH5+
LYVE1 + VEGFR2 + RFP
LYVE1 + VEGFR3 + RFP
C
P16
D
P18

**Supplemental Fig. 3 Analysis of dermal lymphatic surface VEGFR3 at different stages after the induced deletion of *Vegfr2*. A.** Tamoxifen intraperitoneal administration (i.p.) and mice analysis scheme. **B-D.** Analysis of *Vegfr2* deletion in ear skin from the *Vegfr2*$^{iKO;CDH5+}$ and control mice by whole-mount immunostaining for LYVE1 (green) and VEGFR2 (blue, left panel) , and for LYVE1 (green) and VEGFR3 (blue, right panel), at different stage including P14 (B), P16 (C) and P18 (D). The reporter RFP signals indicate the expression of reporter gene (tdTomato) by the Cre (*Cdh5-CreERT2*)-mediated recombination in blood and lymphatic vessels of the *Vegfr2*$^{iKO;CDH5+}$ mice. Scale bar: 200 μm in the left panel of B, C and D; 500 μm in the right panel of B, C and D.

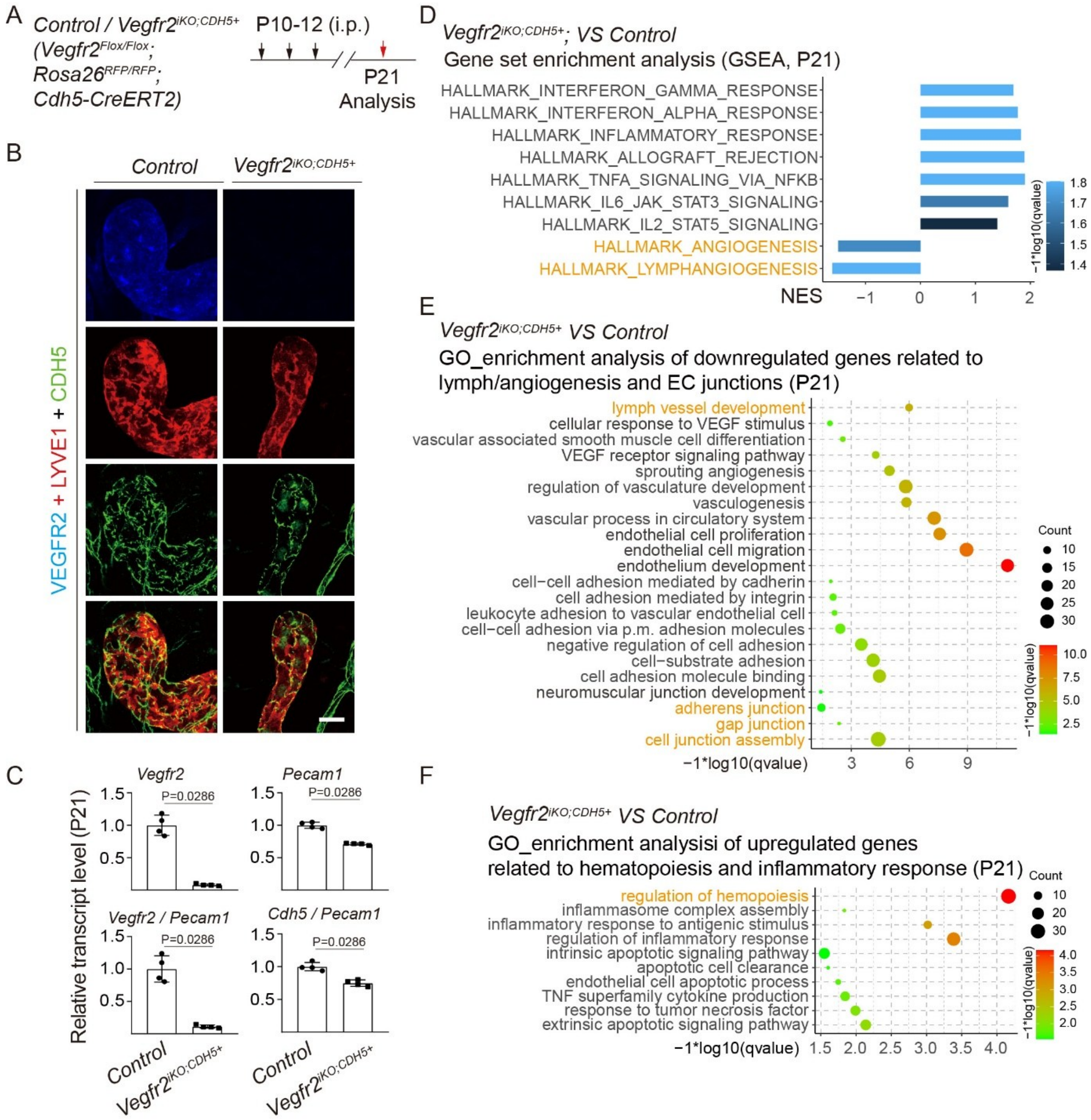


**Supplemental Fig. 4 Alteration of lymphatic morphogenesis and the endothelial transcriptome after the VEGFR2 insufficiency. A.** Tamoxifen intraperitoneal administration (i.p.) and mice analysis scheme. **B.** Analysis of LEC junctions of ear skin from the *Vegfr2*$^{iKO;CDH5+}$ and control mice (P21) by whole-mount immunostaining for LYVE1 (red) and CDH5 (green). *Vegfr2* deletion was confirmed by the whole-mount immunostaining of ear skins for VEGFR2 (red) and CDH5 (green). **C.** Quantification of angiogenic gene transcripts normalized to *Pecam1* and lymphangiogenic gene transcripts normalized to *Lyve1* in ear tissues from the *Vegfr2*$^{iKO;CDH5+}$ and control mice (P21, *Vegfr2*, Control: 1.00 ± 0.16; *Vegfr2*$^{iKO;CDH5+}$: 0.08 ± 0.02, P = 0.0286; *Pecam1*, Control: 1.00 ± 0.05; *Vegfr2*$^{iKO;CDH5+}$: 0.71 ±

0.02, P = 0.0286; *Vegfr2*/*Pecam1*, Control: 1.00 ± 0.20; *Vegfr2*$^{iKO;CDH5+}$: 0.11 ± 0.02, P = 0.0286; *Cdh5*/*Pecam1*, Control: 1.00 ± 0.06; *Vegfr2*$^{iKO;CDH5+}$: 0.75 ± 0.05, P = 0.0286; n = 4 for each group). **D**. GSEA analysis showed the decreased enrichment in lymph/angiogenesis pathways while there was an increased enrichment in genes related to inflammatory responses. **E-F**. GO analysis revealed that the genes related to lymph/angiogenic morphogenesis were significantly downregulated, including endothelial cell migration and proliferation, cell-cell and cell-matrix adhesion, and cell junctions (**E**), while those related to hematopoiesis, endothelial apoptosis and inflammatory responses were significantly upregulated (**F**). Scale bar: 20 μm in B.

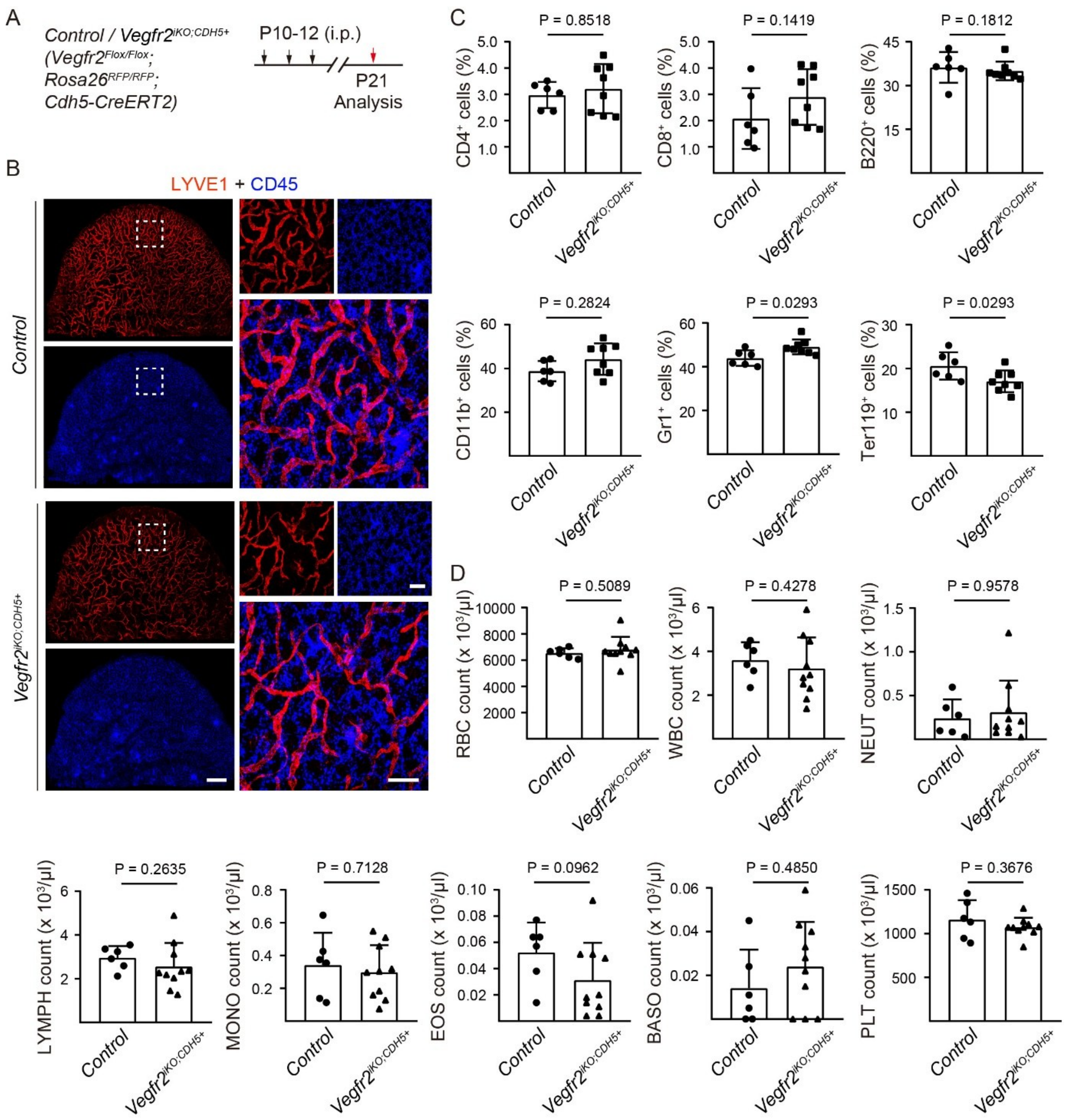


**Supplemental Fig. 5 Analysis of local or systemic immune cells upon VEGFR2 insufficiency. A.** Tamoxifen intraperitoneal administration (i.p.) and mice analysis scheme. **B.** Whole-mount immunostaining of lymphatic vessels in the ear skin from the *Vegfr2*$^{iKO;CDH5+}$ and control mice (P21) for LYVE1 (green) and CD45 (blue). **C.** Flow cytometry analysis for the percentage of immune cells in the bone marrow samples from the *Vegfr2*$^{iKO;CDH5+}$ and control mice (P21; CD4$^+$ cells, Control: 2.97% ± 0.50%, *Vegfr2*$^{iKO;CDH5+}$: 3.21% ± 0.94%, P = 0.8518; CD8$^+$ cells, Control: 2.08% ± 1.16%, *Vegfr2*$^{iKO;CDH5+}$: 2.90% ± 1.06%, P = 0.1419; B220$^+$ cells, Control: 36.22% ± 5.27%, *Vegfr2*$^{iKO;CDH5+}$: 35.02% ± 3.19%, P = 0.1812; CD11b$^+$ cells, Control: 38.79% ± 4.65%, *Vegfr2*$^{iKO;CDH5+}$: 44.29% ± 7.20%, P = 0.2824; Gr1$^+$ cells, Control: 43.92% ±

3.54%, *Vegfr2*$^{iKO;CDH5+}$: 49.03% ± 3.38%, P = 0.0293; Ter119$^{+}$ cells, Control: 20.56% ± 3.13%, *Vegfr2*$^{iKO;CDH5+}$: 17.06% ± 2.48%, P = 0.0293; n = 6 for Control group, n = 8 for *Vegfr2*$^{iKO;CDH5+}$ group). **D.** Analysis of peripheral blood from *Vegfr2*$^{iKO;CDH5+}$ and control mice (X $10^3$/µl; red blood cells-RBC: Control= 6543.33 ± 363.30, *Vegfr2*$^{iKO;CDH5+}$= 6803.00 ± 971.01, P = 0.5089; white blood cells-WBC: Control= 3.61 ± 0.81, *Vegfr2*$^{iKO;CDH5+}$= 3.22 ± 1.42, P = 0.4278; neutrophils-NEUT: Control= 0.24 ± 0.22, *Vegfr2*$^{iKO;CDH5+}$= 0.31 ± 0.36, P = 0.9578; lymphocytes-LYMPH: Control= 2.96 ± 0.53, *Vegfr2*$^{iKO;CDH5+}$= 2.56 ± 1.08, P = 0.2635; monocytes-MONO: Control= 0.34 ± 0.20, *Vegfr2*$^{iKO;CDH5+}$= 0.30 ± 0.17, P = 0.7128; eosinophils-EOS: Control= 0.05 ± 0.02, *Vegfr2*$^{iKO;CDH5+}$= 0.03 ± 0.03, P = 0.0962; basophils-BASO: Control= 0.01 ± 0.02, *Vegfr2*$^{iKO;CDH5+}$= 0.02 ± 0.02, P = 0.4850; platelets-PLT: Control= 1159.67 ± 221.92, *Vegfr2*$^{iKO;CDH5+}$= 1073.40 ± 108.72, P = 0.3676; n = 6 for Control group, n = 10 for *Vegfr2*$^{iKO;CDH5+}$ group). Scale bar: 1000 µm (left panel) and 200 µm (right panel) in B**.**

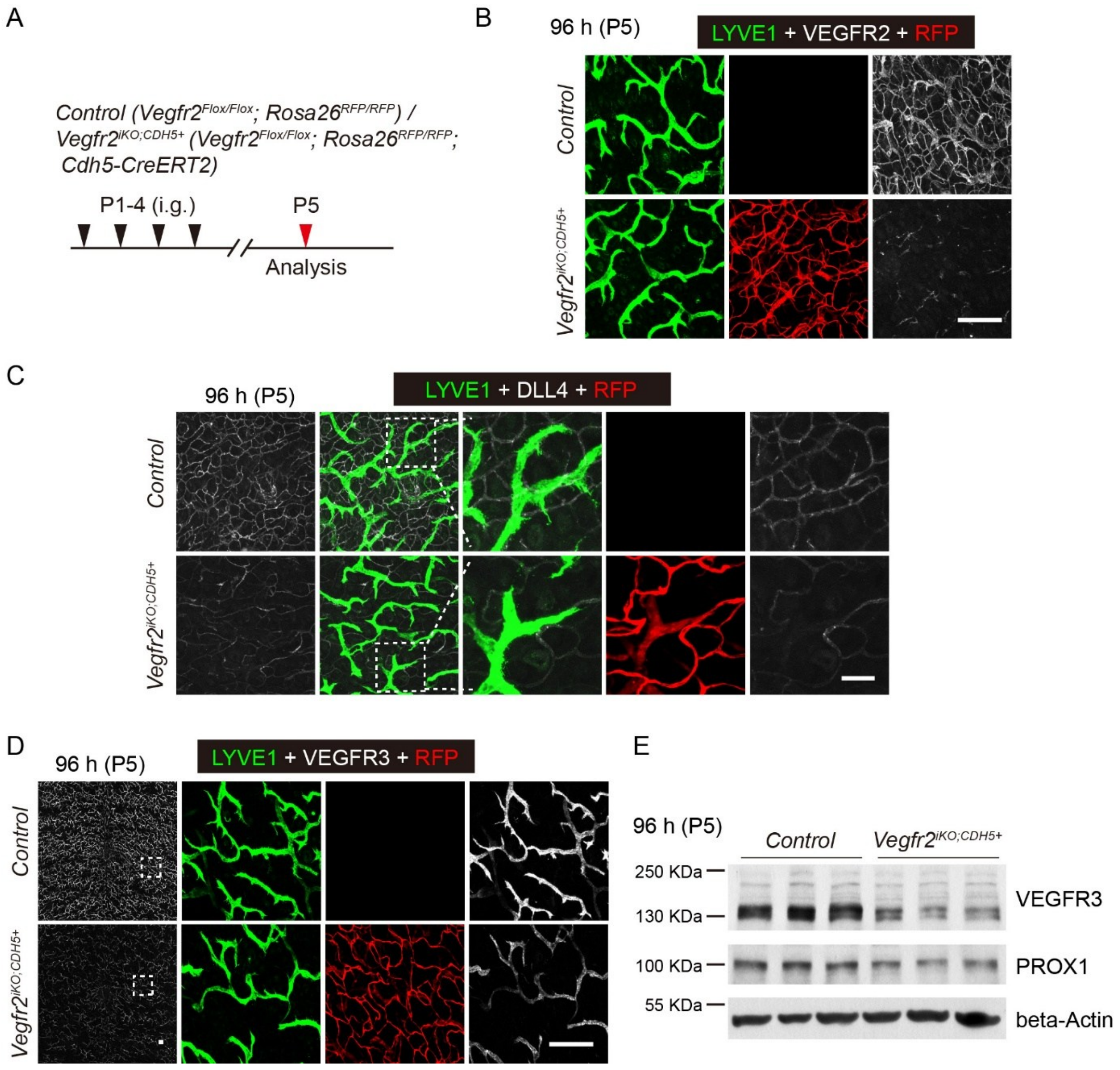


**Supplemental Fig. 6 Alteration of the dermal lymphatic VEGFR3 expression 96 hours after the endothelial VEGFR2 insufficiency. A.** Tamoxifen intragastric administration (i.g.) and mice analysis scheme. **B-D.** Whole-mount immunostaining of the abdominal skin from the *Vegfr2*$^{iKO;CDH5+}$ and control mice (P5) for LYVE1 (green), VEGFR2 (B, white), DLL4 (C, white) or VEGFR3 (D, white). **E.** Western blotting analysis VEGFR3 and PROX1 protein in back skin from the *Vegfr2*$^{iKO;CDH5+}$ and control mice (P5). The reporter RFP signals indicate the Cre (*Cdh5-CreERT2*)-mediated recombination in blood vascular and lymphatic vessels of the *Vegfr2*$^{iKO;CDH5+}$ mice. Scale bar: 200 μm in B and D; 50 μm in C.

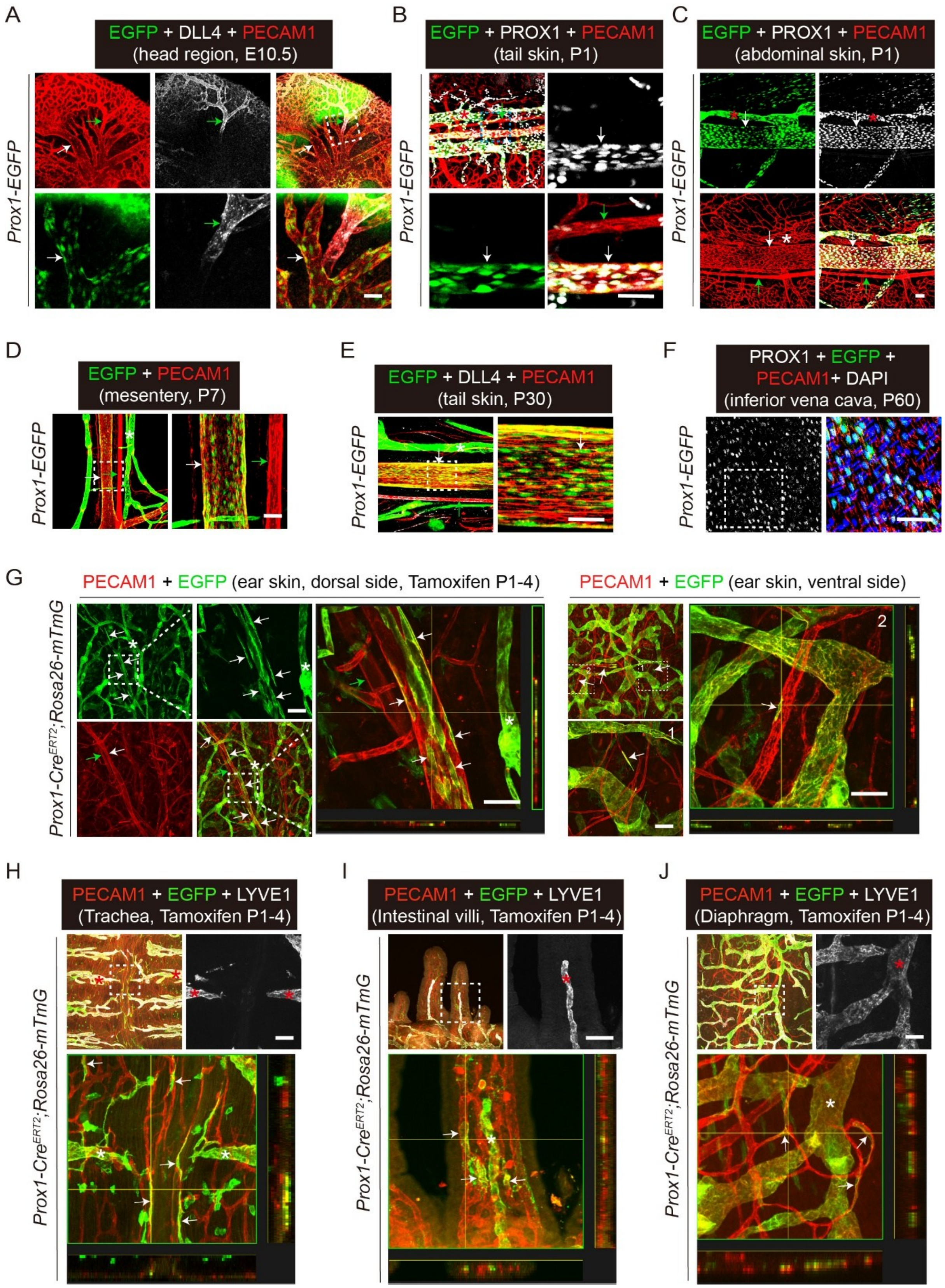


**Supplemental Fig. 7 Detection of PROX1 expression in a proportion of blood vascular endothelial cells in veins and capillaries. A-F.** Whole-mount immunostaining for PECAM1 (red), DLL4 (white) or PROX1 (white) at different developmental stages of various tissues including head (A, E10.5), tail skin (B, P1; E, P30), abdominal skin (C, P1), mesentery (E, P7) and inferior vena cava (F, P60) from *Prox1-EGFP* mice. The reporter EGFP signals indicate PROX1$^{+}$ cells in the *Prox1-EGFP* mice. **G-J.** Whole-mount immunostaining for PECAM1 (red) and LYVE1 (white) of various tissues including skin (G), trachea (H), intestinal villi (I) and diaphragm (J) from *Prox1-CreERT2;Rosa-mTmG* mice at P7. The reporter EGFP signals indicate PROX1$^{+}$ cells in the *Prox1-CreERT2;Rosa-mTmG* mice after tamoxifen treatment. White arrows indicate veins, green arrows indicate arteries, red asterisks indicate lymphatic vessels. Scale bar: 50 μm in A-J.

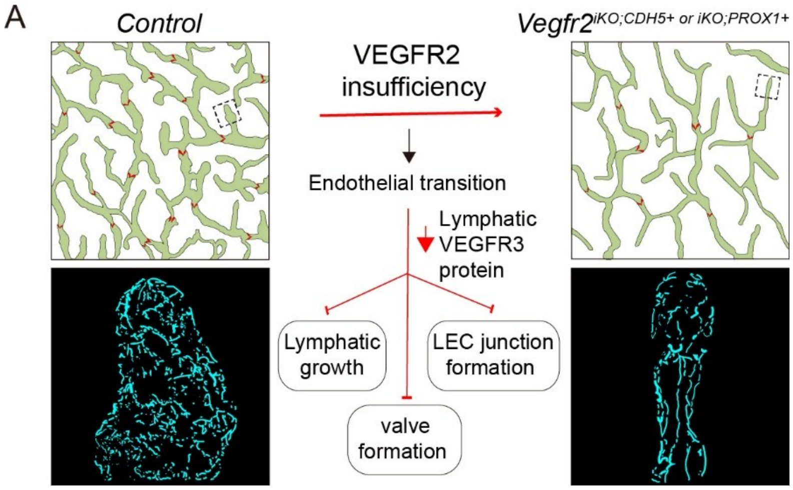


**Supplemental Fig. 8 Schematic illustration of endothelial VEGFR2 in the regulation of lymphatic morphogenesis**. VEGFR2 participates in the regulation of lymphatic development via VEGFR3, a key regulator mediating lymphatic sprouting, valve morphogenesis and LEC button junction formation. Based on the findings from this study employing the endothelial and lymphatic *Vegfr2* mutant mice as well as the *in vitro* LEC work, we show that VEGFR2 is required for the maintenance of the LEC identity and that its insufficiency may trigger a transcriptional trend of LEC to hematopoietic transition.
